# Synaptic histamine shapes the neurocomputational dynamics of human learning

**DOI:** 10.1101/2025.10.05.680501

**Authors:** Michael J. Colwell, Fin van Uum, Philip J. Cowen, Marieke A.G. Martens, Michael Browning, Helen C. Barron, Catherine J. Harmer, Susannah E. Murphy

## Abstract

Histamine was the first canonical monoamine identified in the mammalian brain ^1–6^, yet arguably remains the least understood in its mechanistic contributions to human behaviour. Using a first-in-class causal probe (H_3_R inverse agonist pitolisant), we show how increased synaptic histamine shapes offline and online temporal–hippocampal dynamics, sustaining learning-related activity and polarising retrieval computations. At a broader level, elevated histamine adaptively shifts neurocomputational strategy under high cognitive load, while stabilising value updates during aversive reinforcement learning. These findings uncover a mechanistically grounded influence of this underexplored system on human neurocomputation, supporting its therapeutic potential in psychiatry.

## Introduction

Histamine was observed in the cortex nearly a century ago ^1^, yet its contribution to human behaviour remains elusive compared to classical counterparts such as dopamine and serotonin. Originating from its sole nucleus (tuberomammillary nucleus [TMN]), histamine neurons project brain-wide with greater density than well-established systems such as noradrenaline and oxytocin ^7–12^. Notably, a high proportion of histaminergic projections terminate along a conserved cortico-hippocampal pathway including the mammillary bodies – a classical structure long recognised as an essential relay for human memory ^13–18^. Across this pathway, histamine neurons are controlled by their sole autoreceptor, the 3^rd^ histamine receptor (H_3_R; short isoform) ^11,19–22^, expressed almost exclusively in the brain ^23,24^. Its expression peaks in subcortical circuits, where it shapes synaptic plasticity and network dynamics in regions central to memory in preclinical models ^18,25–33^. In humans, however, the causal role of histamine in temporal–hippocampal dynamics and learning remains unclear.

Animal studies support a central role for histamine and H_3_R in memory: endogenous histamine and H_3_R blockade enhance episodic and working memory ^10,32,34,35^, while targeted TMN lesions and H_3_R opto-stimulation increase aversive learning ^36–39^. In humans, by contrast, evidence is sparse: preliminary PET work links H_3_R density in dorsolateral prefrontal cortex (DLPFC) with working memory ^20^, and attempts to causally manipulate histamine yield non-specific neural and behavioural effects in memory paradigms ^32,40^, while its role in aversive learning remains unexplored. This translational gap likely reflects the historic absence of approved methods to raise central histamine *in vivo*, such as blood–brain barrier–permeable H_3_R inverse agonists with greater selectivity than historical probes (*e.g.*, betahistine) ^41,42^. As the therapeutic potential of H_3_R emerges further – including its preclinical anxiolytic and antidepressant properties ^43–45^ – it is becoming increasingly important to understand the broader contribution of histamine to human behaviour.

This longstanding gap may be overcome with pitolisant, a first-in-class H_3_R inverse agonist recently approved for narcolepsy treatment ^46,47^. Notably, pitolisant is an order of magnitude more selective for H_3_R than historical compounds (*e.g.*, > 200-fold selectivity vs. betahistine) ^41,42,48,49^, with preferential activity at H_3_ autoreceptors ^19–22^. Pitolisant produces rapid elevations in synaptic histamine and achieves high forebrain H_3_R occupancy in humans (>80%) ^42,48,50,51^, providing a unique opportunity to probe histamine’s causal role in human behaviour. We hypothesised that pitolisant would reflect preclinical signatures of elevated histamine, shaping temporal–hippocampal circuitry to support human learning and adaptive cognition.

Here, we leverage pharmaco-fMRI to causally reveal how synaptic histamine shapes the neurocomputational dynamics of human learning. This approach captures its influence across offline and online temporal-hippocampal learning states, and places them in broader computational contexts of working memory and reinforcement learning. In line with our predictions, we show elevated synaptic histamine shifts temporal-hippocampal network dynamics, while sustaining learning-related activity and driving divergent retrieval computations. At a broader level, elevated histamine adaptively shifts neurocomputational strategy under high cognitive load, while stabilising value updates during aversive reinforcement learning.

## Results

### Synaptic histamine shapes offline temporal-hippocampal network dynamics

To examine how increased synaptic histamine (via pitolisant) influences the neural dynamics of both offline and online memory processing, we implemented a multi-stage task paradigm (Fig. 1A). In stage one, participants iteratively encoded images until reaching criterion (see Supplementary Note 1). Stage two comprised rsfMRI to assess how synaptic histamine modulates offline temporal-hippocampal network dynamics. In stage three, participants encoded both novel and previously learned (‘familiar’) images during fMRI, enabling a novel > familiar contrast of new learning. Finally, in stage four, recognition memory for images learned throughout the paradigm was probed.

**Fig. 1:**
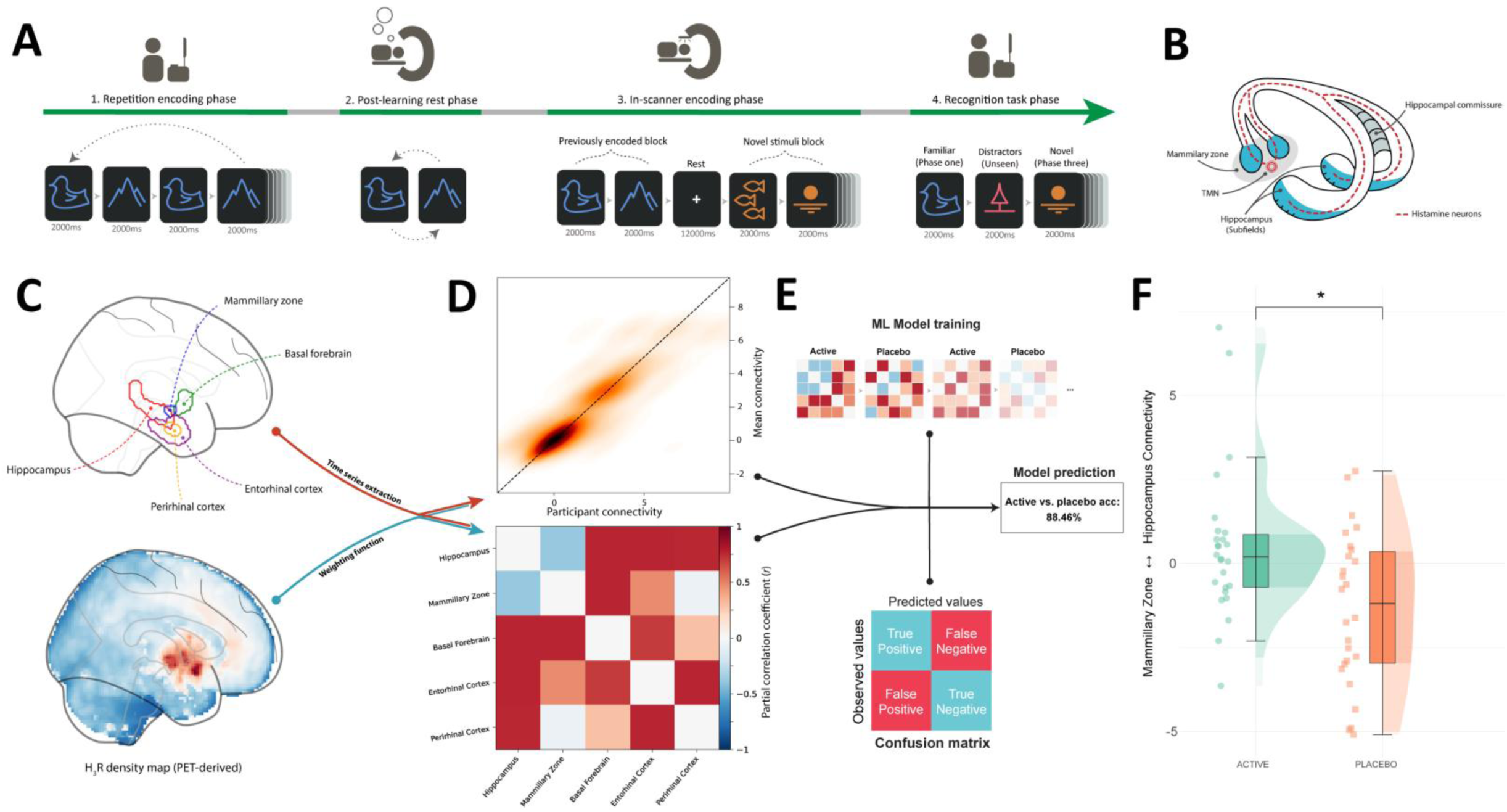
Multi-stage memory paradigm flow and modelling of H3R-weighted network dynamics during post-learning rest. **A** The paradigm comprised four stages: (1) encoding of visual pictures (landscapes/animals) through repeated presentation; (2) post-learning rest (10.4 mins rsfMRI) to capture network dynamics following encoding; (3) further encoding of novel (*n* = 48) and previously learned (*n* = 8) images during fMRI, enabling the novel > familiar contrast; and (4) recognition testing of previously learned (*n* = 56) and novel distractor (*n* = 27) images. **B** Histamine neurons are densely distributed along a pathway critical for memory consolidation and encoding, spanning the mammillary zone (including the TMN) to the hippocampus via the fornix ^13,23^. **C** During post-learning rest (stage two), signal was extracted from memory-relevant ROIs with evidence of H3R sensitivity. Signals were weighted by regional H3R density derived from PET maps ^60^. **D** Weighted signals were assembled into individual covariance matrices and **E** classified via linear discriminant analysis. Cross-validated models distinguished pitolisant from placebo with 88.5% accuracy. **F** Univariate analysis showed that network changes were driven by stronger mammillary zone ↔ hippocampus connectivity in the pitolisant group (permutation testing [pitolisant > placebo]: *t*[51] = 2.92, *p* = 0.0266, FWE-corrected, Cohen’s *d* = 0.81). Boxplots depict the interquartile range (IQR, central line = median), whiskers = ±1.5 × IQR, and half-violins show data distribution. \**p* ≤ 0.05, FWE-corrected. **D**, **F** include data for *N* = 52 individuals. FWE Family-wise error; ML Machine Learning.

As temporal-hippocampal network connectivity is shaped by recent learning ^52^, we investigated if synaptic histamine influences offline network dynamics post-learning (stage two). Signal estimates from memory-relevant regions sensitive to H_3_R manipulation ^13,31,32,39,53–59^ were weighted by global H_3_R density (via *neuromaps* ^60^) and used to construct covariance-based network models (Fig. 1B–D). A cross-validated machine learning classifier distinguished pitolisant vs. placebo with 88.5% accuracy (Fig. 1E). This network-level separation was explained by changes in connectivity between the hippocampus and mammillary zone – a zone containing the origin of histamine neurons ^61,62^ – which increased under pitolisant (Fig. 1F). Results without H_3_R-weighting showed similar group differences, with reduced effect size (Supplementary Note 2).

Together, these findings suggest that synaptic histamine shifts offline temporal-hippocampal dynamics, providing network-level context for subsequent learning, as explored in the following section (stage three; see Fig. 2H–I).

**Fig. 2:**
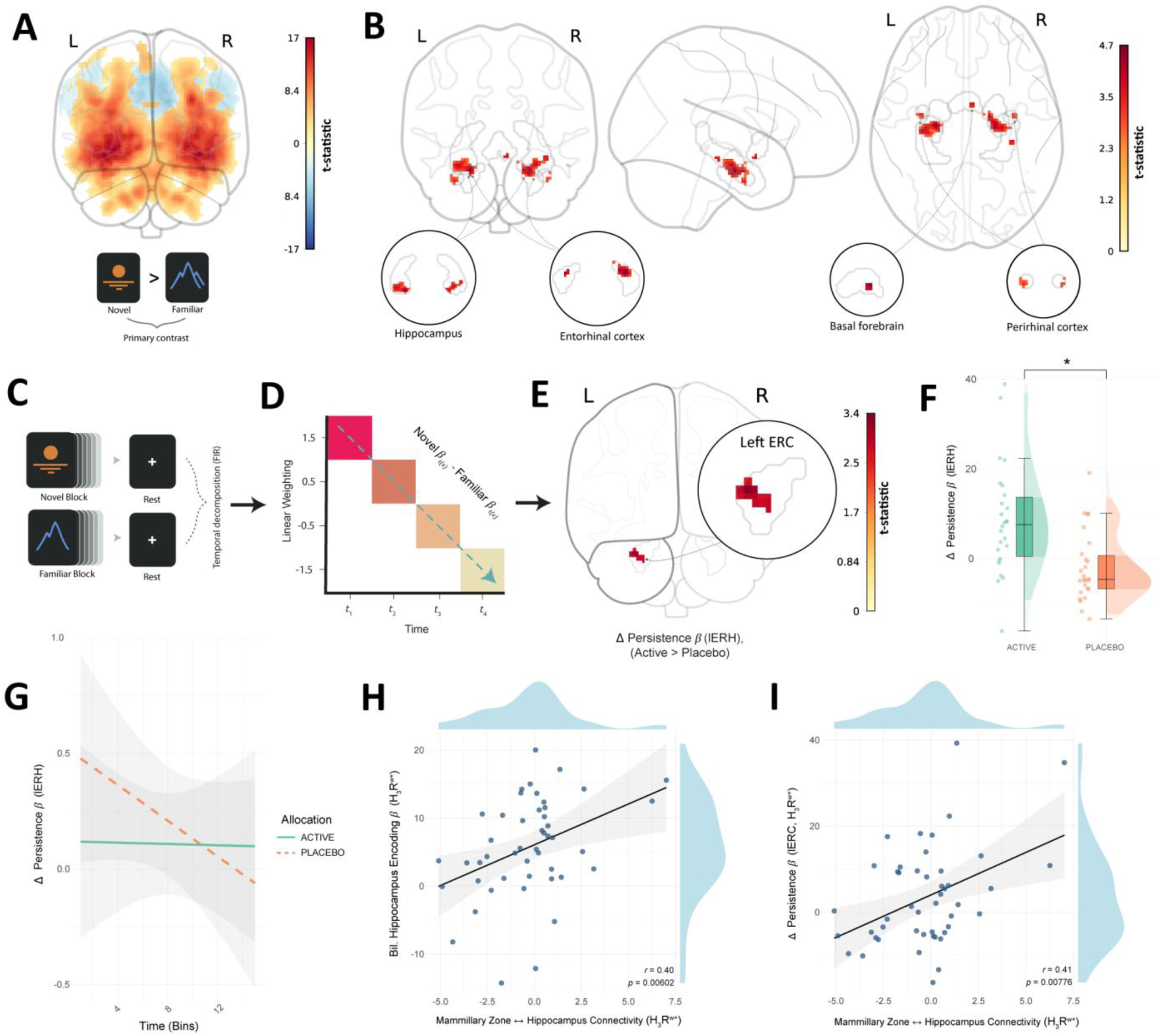
Neural dynamics of new learning and post-encoding signal persistence. During stage three of the memory paradigm, participants encoded novel images and previously learned images in separate blocks (*N*=12 interleaved blocks), separated by post-learning rests (12,000ms). **A** This panel depicts activity specific to new learning (novel > familiar contrast, TFCE-corrected; see Supplementary Note 3 for full statistics and coordinates). **B** Pitolisant increased activation during new learning across an *a priori* memory network, including bilateral hippocampal subfields, basal forebrain, and rhinal cortices (entorhinal, perirhinal; TFCE-corrected SVC). ROI outlines are shown in grey. **C**–**D** Rest periods were temporally modelled with finite impulse response functions to capture signal persistence after new learning (novel > familiar). **E**–**F** Pitolisant sustained signal persistence in the left medial entorhinal cortex (SVC permutation testing: peak MNI = -28, -12, -30; cluster = 19 voxels; *t*(51) = 3.37, *p* = 0.0222, TFCE-corrected). Boxplots depict the interquartile range (IQR, central line = median), whiskers = ±1.5 × IQR, and half-violins show data distribution. \**p* ≤ 0.05, FWE-corrected. **G** Depicts the temporal trajectory (Δ Persistence *β*) of signal persistence for novel > familiar activity during post-encoding rest; regression lines reflect linear fits ± SEM over time. **A**, **B**, **E** – **G** contain data for *N* = 52 individuals. **H**–**I** Hippocampal–mammillary zone connectivity during the prior rest (stage two) predicted both hippocampal encoding activity (β = 1.13, *p* = 0.00602, FWE-corrected; ηp^2^ = 0.16 [95% CI 0.01, 0.35]; *r* = 0.40) and entorhinal signal persistence (β = 1.98, *p* = 0.00422, FWE-corrected; ηp^2^ = 0.19 [0.03, 0.39]; *r* = 0.44). The positive associations in panels **G**–**H** were observable within allocation groups, via aggregated posterior estimates (hippocampus encoding β = 0.607, 95% credible interval [0.606, 0.608]; Δ signal decay β = 1.19 [0.032, 2.36). Scatterplot points depict participant mean datapoints; regression lines reflect linear fits ± SEM. Panels **H**, **I** contain data for *N* = 47 individuals. Abbreviations: BOLD Blood Oxygenation Level Dependent; H3R^W+^ = Histamine 3 receptor weighted; lERC Left entorhinal cortex; ROI Region of Interest; TFCE Threshold-Free Cluster Enhancement.

### Histamine stabilises new learning signal persistence and leads to asymmetrical retrieval computations

Next, we investigated how increased synaptic histamine influences the neural dynamics of new learning. This was tested during the third stage of the multi-stage memory paradigm, where participants encoded novel images alongside those previously learned (‘familiar’) in stage one. Neural activity was contrasted between these conditions (novel > familiar; Fig. 2A; Supplementary Table 3), allowing the isolation of neural dynamics specific to new learning. During new learning, pitolisant increased activation in bilateral hippocampal subfields in both whole-brain and small-volume corrected (SVC) analyses (Fig. 2B; Supplementary Note 3; Supplementary Table 4). Increased activity was also observed across the *a priori* memory network from the preceding rest stage, including basal forebrain, entorhinal cortex and perirhinal cortex. Stimulus discrimination behaviour (accuracy and response time) did not differ significantly between groups, indicating comparable task engagement (Supplementary Note 4).

As persisting temporal-hippocampal activity during post-learning rest is linked to effective consolidation in humans ^63–65^, we asked whether synaptic histamine influences this process. To test this, signal from post-encoding rest intervals was temporally decomposed using finite impulse response (FIR) modelling, enabling us to capture signal persistence after new learning (novel > familiar; Fig. 2C–D). Pitolisant sustained temporal persistence for new learning in the left medial entorhinal cortex (Fig. 2E–F). This effect reflects persistence beyond initial response magnitude, as shown by the FIR-isolated decay trajectories (Fig. 2G). Importantly, encoding-related activity in the left hippocampus and entorhinal cortex – but not their right-hemisphere homologues – predicted entorhinal persistence, suggesting a lateralised effect (Supplementary Note 5). Moreover, offline hippocampal–mammillary zone connectivity (stage two) was associated with higher hippocampal activity during encoding (Fig. 2H) and increased entorhinal signal persistence (Fig. 2I). These associations were absent or weakened when H_3_R-weighting was removed (Supplementary Table 9), supporting histaminergic specificity.

Thirty minutes after in-scanner encoding (stage three), recognition memory for images learned throughout the paradigm was probed (stage four). Choice behaviour was modelled with a drift diffusion model (DDM) of evidence accumulation. Parameter estimation was stimulus-dependent, as previously encoded images and unseen distractors are assumed to differ in their evidence accumulation properties ^66^. Full details of model selection, parameter recoverability, and posterior predictive checks are provided in the Supplementary Methods. The best-fitting model included four stimulus-dependent parameters, formalising distinct parts of choice behaviour: drift rate (*v*_*s*_), reflecting evidence accumulation efficiency; decision policy (*a*_*s*_), reflecting the evidence threshold required before making a decision, capturing signal-noise separation; non-decision time (*t*_*er*,*s*_), reflecting the pre-decisional buffer between stimulus onset and evidence accumulation; and starting point bias (*z*_*s*_), reflecting an initial bias toward one response boundary.

Behaviourally, pitolisant improved recognition accuracy (mean %; ANOVA main effect of group: F[1,50] = 8.72, *p* = 0.0048, η_p_^2^ = 0.13 [95% CI 0.01, 0.31]) and reduced time to choice (ms; ANOVA main effect of group: F[1,50] = 8.63, *p* = 0.005, η_p_^2^ = 0.13 [0.01, 0.31]) across stimulus types (Fig. 3A–B). Computationally, pitolisant increased drift rate for previously encoded images only (*v*_*s*_ ANOVA group × stimulus interaction: F[1,50] = 2.48, *p* = 0.000973, η_p_^2^ = 0.12 [0.03, 0.25]; *v*_*s*_ ANOVA main effect of group: F[1,50] = 3.68, *p* = 0.06; Fig. 3C). Conversely, pitolisant reduced decision policy for distractors only (log *a*_*s*_ ANOVA group × stimulus interaction: F[1,50] = 6.17, *p* = 0.0164, η_p_^2^ = 0.11 [0.00, 0.29]; log *a*_*s*_ main effect of group: F[1,50] = 0.681, *p* = 0.413; Fig. 3D). No significant group differences emerged for other parameters (Supplementary Fig. 7). Post-encoding signal persistence (lERC; stage three) was positively associated with drift rate for previously encoded images but not distractors (Fig. 3E). Conversely, persistence negatively predicted decision policy for distractors, but not previously encoded items (Fig. 3F).

**Fig. 3:**
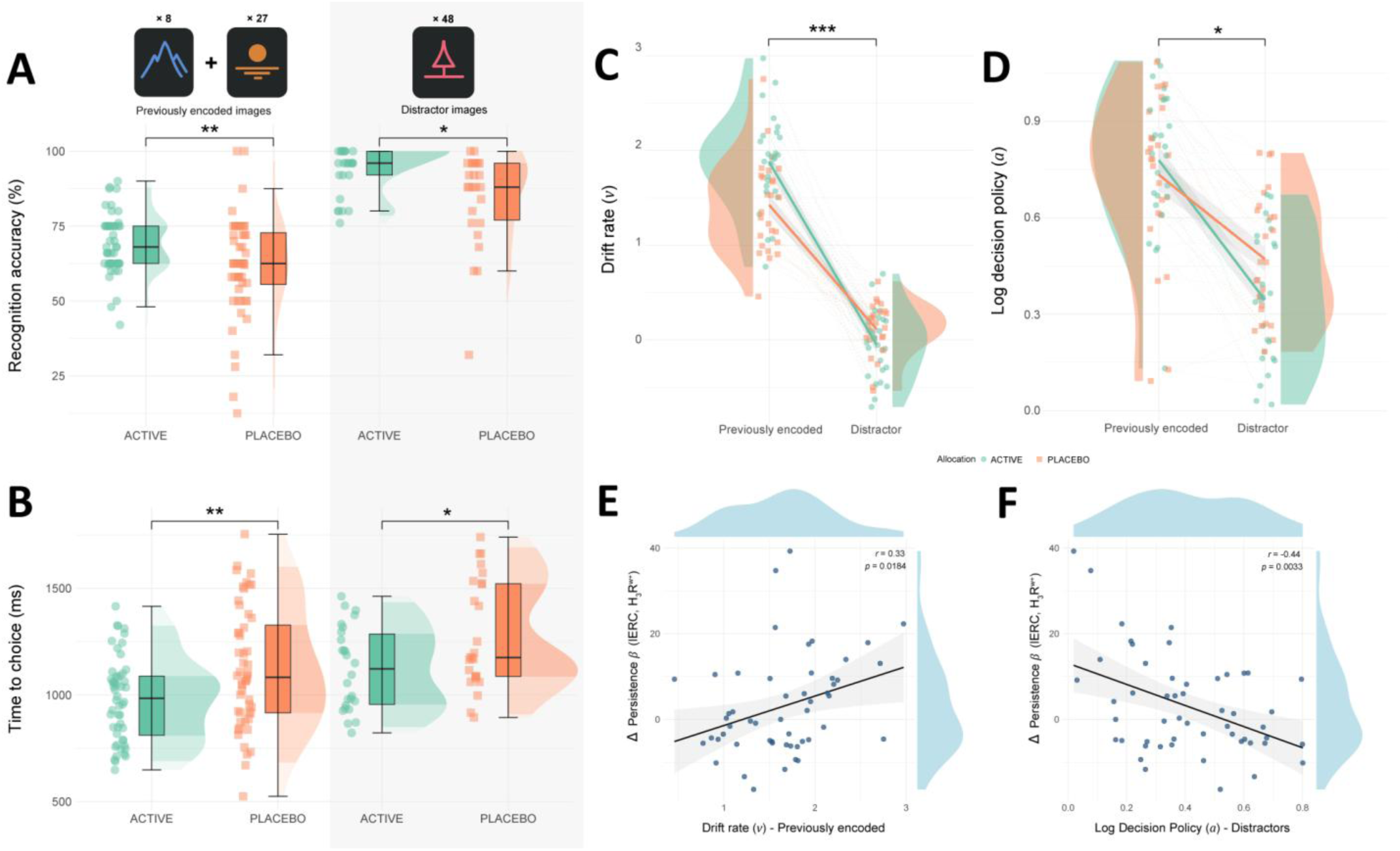
Decision computations during memory recognition and their association with prior encoding-related activity. The multi-stage memory paradigm concluded with a memory recognition task. Test items comprised previously encoded images (stage one: *n*=8; stage three: *n*=27) and unseen distractors (*n*=48). **A** Recognition accuracy was higher under pitolisant (distractors EMM = 8.62 ± 3.71, two-tailed *p* = 0.022, *d* = 0.76 [0.11, 1.40]; previously encoded EMM = 7.50 ± 2.69, two-tailed *p* = 0.014, *d* = 0.67 [0.13, 1.18]). **B** Time to correct choices was reduced under pitolisant (distractors EMM = -154.0 ± 66.0, two-tailed *p* = 0.02, *d* = -0.81 [-1.50, -0.12]; previously encoded EMM = 144.0 ± 54.5, two-tailed *p* = 0.010, *d* = -0.76 [-1.34, -0.18]). **C** Drift rate (***v***_***s***_) was higher under pitolisant for previously encoded images (EMM = 0.45 ± 0.12, *p* = 0.0002, *d* = 1.09) but not distractors (EMM = -0.17 ± 0.12, two-tailed *p* = 0.1562). **D** Decision policy (log ***a***_***s***_) was reduced for distractors under pitolisant (EMM = -0.13 ± 0.06, two-tailed *p* = 0.0384, *d* = 0.72), with no effect for previously encoded items (EMM = 0.05 ± 0.06, two-tailed *p* = 0.456). **E** lERC signal persistence predicted drift rate for previously encoded items (β = 0.02, two-tailed *p* = 0.0368, FWE-corrected; ηp^2^ = 0.11 [0.00, 0.28]; *r* = 0.33), but not for distractors (β = -0.0005, two-tailed *p* = 0.897). **F** lERC persistence negatively predicted decision policy for distractors (β = -0.008, two-tailed *p* = 0.0033, FWE-corrected; ηp^2^ = 0.19 [0.04, 0.38]; *r* = -0.44), but not previously encoded items (β = 0.0005, two-tailed *p* = 0.838). The direction of effects in **E**–**F** was consistent across stimulus types (adjusted for groups), confirmed by posterior estimates (***v***_***s***_β = 0.006 [0.004, 0.008]; log ***a***_***s***_β = -0.004 [-0.007, -0.0006]). **A**–**F** include *N*=52 individuals. **A**–**B** Boxplots depict the interquartile range (IQR, central line = median), whiskers = ±1.5 × IQR, and half-violins show data distribution. **C**–**D** Points/violins show data distribution; bold lines depict group means. *** *p* ≤ 0.001, ** *p* ≤ 0.01, * *p* ≤ 0.05, FWE-corrected. **E**–**F** Scatterplot points depict participant mean datapoints; regression lines reflect linear fits ± SEM. Abbreviations: H3R^W+^ = Histamine 3 receptor weighted; ms milliseconds; lERC Left entorhinal cortex; TFCE Threshold-Free Cluster Enhancement.

Taken together, these results indicate that synaptic histamine enhances entorhinal signal persistence following learning, while during retrieval it drives asymmetrical computations – selectively enhancing evidence accumulation for past learning and lowering the evidence required for unfamiliar information. More broadly, our findings suggest a histaminergic shaping of temporal–hippocampal dynamics across offline and online learning states in support of consolidation.

### Synaptic histamine influences the neurocomputational signatures of working memory

We next extended our investigation from episodic learning to working memory, asking how elevated synaptic histamine shapes its neurocomputational signatures. Participants performed an fMRI-adapted verbal *n*-back task (Fig. 4A), indicating whether a target letter had appeared *n* trials earlier (0-back [control], 1-back [easy], 2-back [moderate], 3-back [hard]). This stepwise design enabled the modelling of load-dependent changes in decision computations and neural activity. As with the memory recognition task, choice behaviour was modelled using a drift diffusion model (DDM; Fig. 4B). The winning model was selected based on goodness-of-fit among candidate models and parameter recoverability (Supplementary Methods), and validated by posterior predictive checks (Fig. 4C). The final model included three parameters: drift rate (+*v*/−*v*), non-decision time (*T*_*er*_), and decision policy (*a*).

**Fig. 4:**
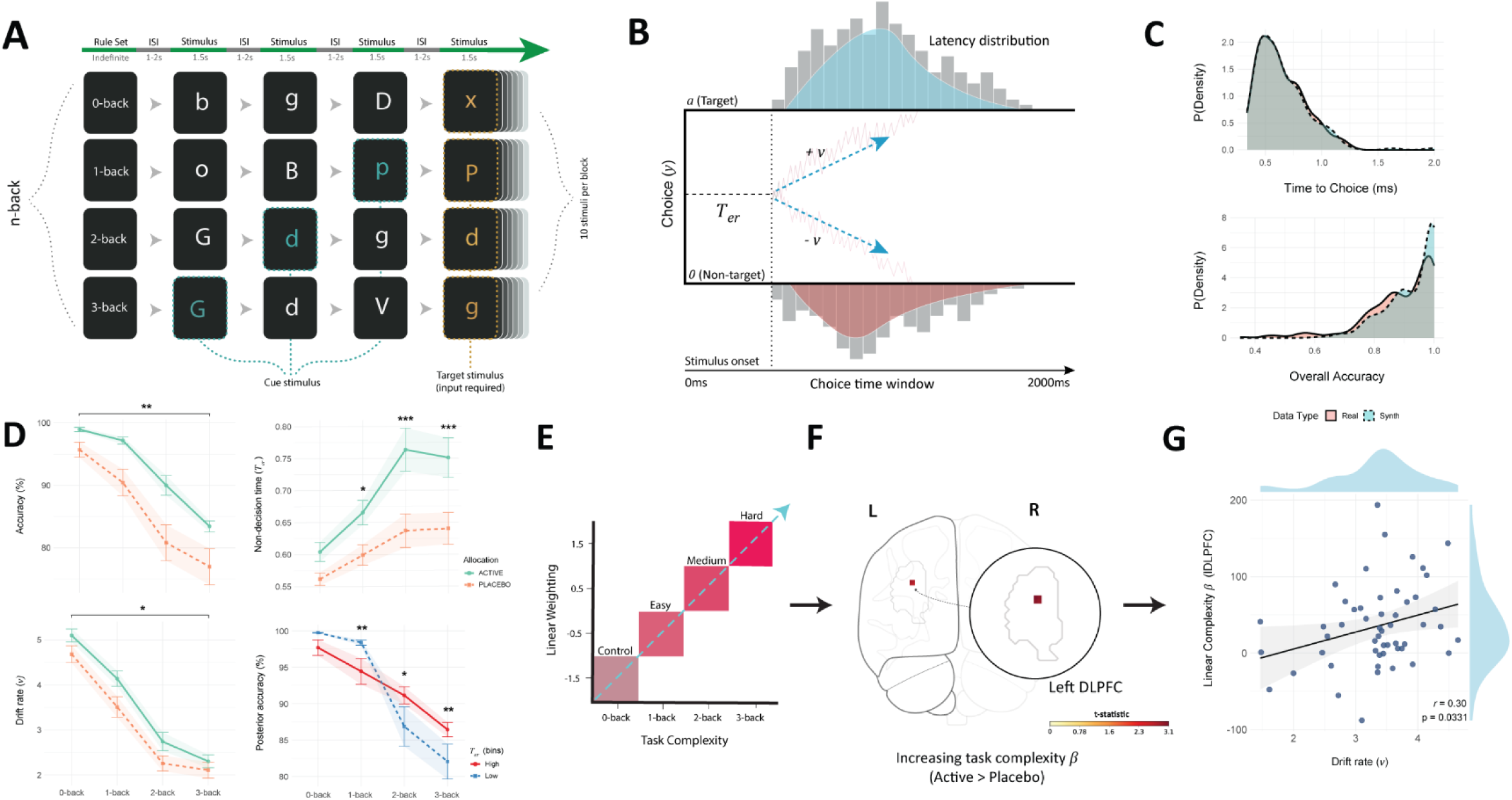
Task procedure, computational modelling and analysis of the complex working memory task. **A** The verbal n-back task included four levels of increasing complexity (0–3 back; rows), with each requiring cued information to be maintained over longer intervals. Before each block, participants were cued with task rules (*e.g.*, “press right if the current letter matches that two trials ago”). Each level comprised four blocks (16 total), with 20,000ms rests interleaved. **B** Drift diffusion models were fit to choice behaviour (4 chains × 4,000 samples per *N*), containing three parameters: drift rate (+*v*/−*v*), non-decision time (*T*_*er*_) and decision policy (*a*). **C** Models fit to individual participants resulted in posteriors which closely matched observed behaviour. **D** Pitolisant increased accuracy (upper left) and drift rate (lower left) independent of task load, while non-decision time increased as a function of load (upper right). Splitting trials by high vs. low *T*_*er*_ (within-subjects median split) revealed that higher *T*_*er*_ decreased accuracy under low load, but improved accuracy at high load (lower right). Plot points depict mean value; error bars and shaded area depict SEM. \*\*\**p* ≤ 0.001, \*\**p* ≤ 0.01, \**p ≤* 0.05 by two-tailed EMM tests, FWE-corrected (see Supplementary Note 6 for full statistics). **E** BOLD signal was estimated as a function of task complexity via parametric modelling. **F** Pitolisant increased recruitment of the left DLPFC as task complexity increased (SVC permutation testing: peak MNI = -34, 38, 34; cluster = 5 voxels; *t*(51) = 3.12, *p* = 0.0456, TFCE-corrected). **G** Linear activity within this cluster predicted drift rate ***v*** (β = 0.004, *p* = 0.0335; ηp^2^ = 0.09 [0.00, 0.26]; *r* = 0.30); this relationship was observable within allocation groups, based on aggregated posterior estimates (β = 0.003 [0.001, 0.005]). Scatterplot points depict participant mean datapoints; regression lines reflect linear fits ± SEM. Abbreviations: lDLPFC = Left Dorsolateral Prefrontal Cortex; H3R^W+^ = Histamine 3 receptor weighted; TFCE = Threshold-Free Cluster Enhancement.

Behaviourally, pitolisant improved overall task accuracy (main effect of group ANOVA: F[1,50] = 7.89, *p* = 0.0070, η_p_^2^ = 0.14 [0.01, 0.32]; Fig. 4D, upper left), but did not significantly change time to choice (Supplementary Fig. 9A). Computationally, pitolisant increased drift rate (+*v*/−*v*) independent of task load (main effect of group ANOVA: F[1,50] = 4.94, *p* = 0.0308, η_p_^2^ = 0.09 [0.00, 0.26]; Fig. 4D, lower left), while no significant group effect on decision policy (*a*) was observed (Supplementary Fig. 9B). Non-decision time (*T*_*er*_) increased with task complexity under pitolisant (ANOVA group × task condition interaction: F[3,150] = 2.70, *p* = 0.0481, η_p_ ^2^ = 0.05 [0.00, 0.11]; Fig. 4D, upper right). To assess its contribution to behaviour, we split posterior performance into high and low *T*_*er*_ bins (500,000 draws; see Supplementary Methods). High *T*_*er*_ reduced accuracy at low load but enhanced it at high load (ANOVA group × task condition interaction: F[3, 158] = 2.78, p = 0.043, η_p_^2^ = 0.08 [0.01, 0.15]; Fig. 4D, lower right), consistent with confirmatory analyses of observed accuracy (Supplementary Fig. 9C; Supplementary Note 7). Under higher working-memory load, encoding demands additional resources ^67^, and increases in *T*_*er*_ may reflect a strategic reallocation of pre-decisional time to support these processes. Importantly, such slowing of *T*_*er*_ can reflect adaptive shifts in cognitive strategy, independent of visuomotor deadtime ^68^.

In a parametric model capturing neural responses to increasing task load (Fig. 4E), pitolisant increased recruitment of the left dorsolateral prefrontal cortex (Fig. 4F; Supplementary Fig. 11). Load-related activity in this region tracked drift rate (*v*; Fig. 4G). Notably, these effects were absent when H_3_R-weighting was removed from parameter estimates (Supplementary Table 9), supporting histaminergic specificity. Furthermore, drug-related increases in neural recruitment peaked at moderate task load (2 > 0-back), where DLPFC activity plateaus (Supplementary Fig. 12), encompassing left DLPFC, bilateral hippocampus, and substantia nigra (in both whole-brain and SVC analyses; Supplementary Figs. 10–11, 13; Supplementary Tables 6–7).

Overall, these findings indicate that elevated synaptic histamine promotes adaptive shifts in neurocomputational strategy under high cognitive loads. Specifically, the interval preceding evidence accumulation lengthens (*T*_*er*_), while recruitment of the left dorsolateral prefrontal cortex increases – a region which tracks evidence accumulation efficiency (*v*), as seen here and in previous causal interference work ^69,70^.

### H3R blockade influences aversive computations during reinforcement learning

Next, we investigated whether increasing synaptic histamine influences reinforcement learning using a probabilistic instrumental learning task (Fig. 5A). Throughout the task, participants chose between symbol pairs associated with stable outcome probabilities, in either rewarding (monetary gain) or aversive (monetary loss) contexts. Optimal performance required selecting the symbol with the higher probability (70%) of favourable outcome (winning money or avoiding loss). Participant choices throughout the task were fit to a computational Q-learning model (Fig. 5B) adapted from ^71–73^(see Supplementary Methods for model specifications and validation procedures). The model formalises learning and decision computations with two parameters: learning rate (*⍺*) which describes the rate at which expectations are updated during learning; and inverse decision temperature (*β*), which captures the degree of stochasticity during decision-making. As *⍺* and *β* covary, inferential analyses incorporated the reciprocal parameter in the linear model structure (see Supplementary Methods for further details).

**Fig. 5:**
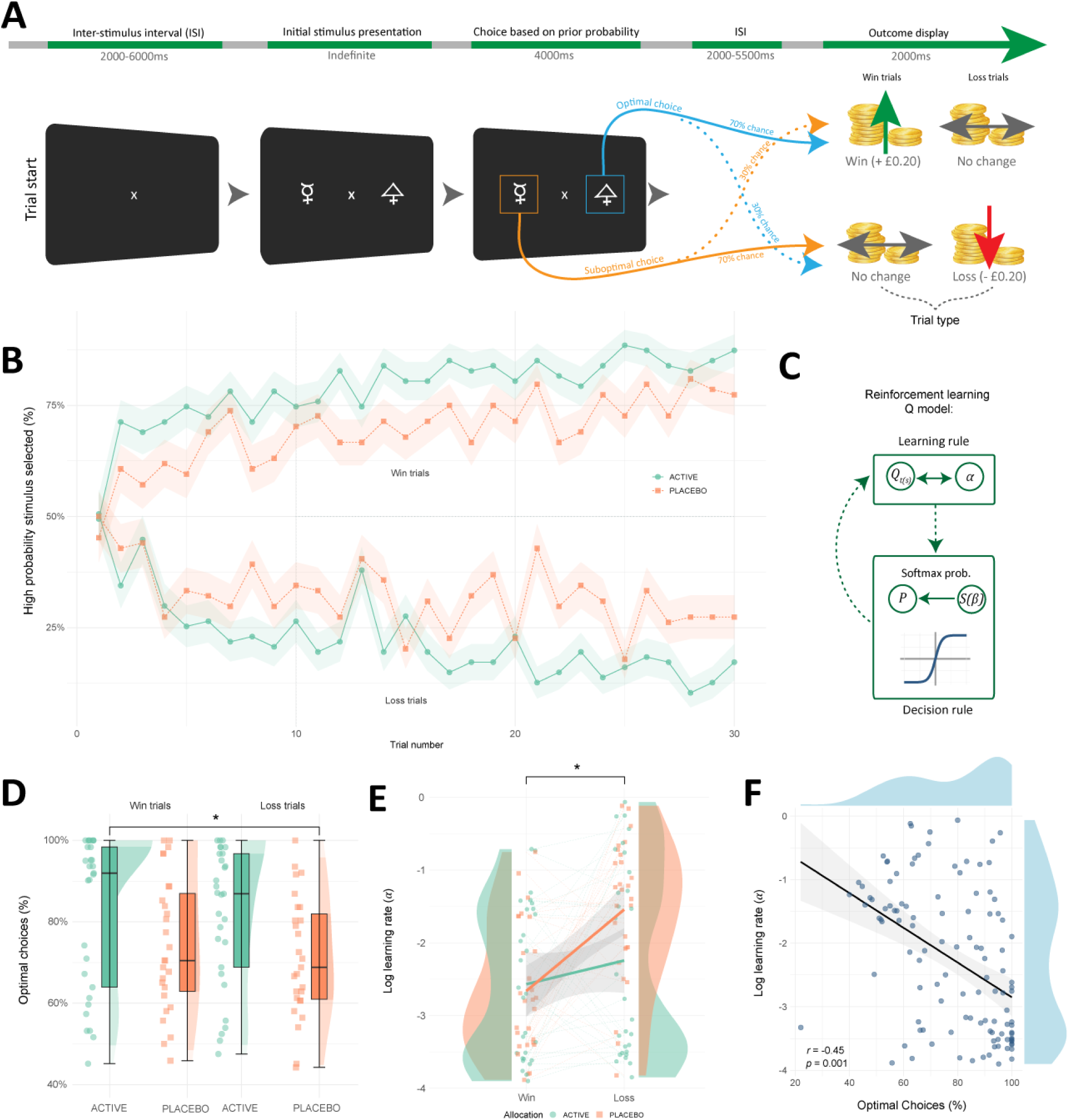
Reinforcement learning procedure, computational modelling and causal analyses. **A** Probabilistic instrumental learning task. Participants chose between symbols within a pair, with two interleaved pairs representing win and loss conditions. Symbols had fixed reciprocal probabilities (70% vs. 30%) of favourable outcomes (winning or avoiding loss). Learning occurred across 60 trials (30 per condition) over three blocks. Optimal performance required repeated selection of the high-probability symbol; cumulative earnings were paid at study completion. **B** Learning over time across groups (averaged across blocks). The Y-axis (‘high probability stimulus selected’) refers to the selection (mean %) of high probability win or loss symbols. **C** Choice behaviour was fit to a Q-learning model ^71–73^, comprising a learning rule which updates value estimates (*Q*_*t*(*s*)_), and decision rule assigning choice probability (*P*) via softmax. Two parameters, learning rate (*⍺*) and inverse decision temperature (*β*), formalise distinct computational processes contributing to task behaviour (for further details, see Supplementary Methods). **D** Pitolisant increased overall optimal choices (win trials EMM = 9.57 ± 4.62, *p* = 0.041, *d* = 0.74 [0.23, 1.46]; loss trials EMM = 11.37 ± 4.62, *p* = 0.016, *d* = 0.88 [0.16, 1.60]). Boxplots depict the interquartile range (IQR, central line = median), whiskers = ±1.5 × IQR, and half-violins show data distribution. **E** Pitolisant reduced learning rate (log *⍺*) during loss trials (win trials EMM = 0.22 ± 0.30, *p* = 0.74; loss trials EMM = -0.56 ± 0.26, two-tailed *p* = 0.0364, *d* = -0.59 [-1.15, -0.03]). Points/violins show data distribution; bold lines depict group means. **F** Learning rate (log *⍺*) was negatively associated with optimal choices (β = -4.32, two-tailed *p* = 2.31e-05, ηp^2^ = 0.16 [0.05, 0.29); *r* = -0.45); this relationship was observable within allocation groups, based on aggregated posterior estimates (β = -6.70 [-7.13, -6.26]). Scatterplot points depict participant mean datapoints; regression lines reflect linear fits ± SEM. Panels **C**–**F** contain data from *N* = 57 individuals. * *p* ≤ 0.05 by two-tailed estimated marginal means tests (FWE-corrected). ISI Inter-stimulus interval.

Throughout the task, pitolisant increased optimal choices (Fig. 5C–D; main effect of group ANOVA: F[1,55] = 7.08 *p* = 0.0102, η_p_^2^ = 0.11 [0.01, 0.28]), but did not significantly change time to choice (Supplementary Fig. 14A). Computational modelling showed that pitolisant reduced the learning rate *⍺* during loss trials (Fig. 5E; ANOVA group × task condition interaction: F[1,55] = 4.29, *p* = 0.041963, η_p_^2^ = 0.06 [0.00, 0.19]; main effect of group ANOVA: F[1,55] = 0.68, *p* = 0.413), but did not significantly change inverse decision temperature *β* (Supplementary Fig. 14B; Supplementary Note 8). Moreover, these effects persisted when the first ten trials per condition were excluded from the computational model, indicating they were not driven by early exploratory noise (Supplementary Note 8). Notably, both observed optimal choices and model-derived (posterior) optimal choices were associated with lower *⍺* (Fig. 5F; Supplementary Fig. 14C); consistent with the principle that, in stable environments, lower *⍺* is optimal given that prior outcomes are more predictive of future uncertainty ^74,75^. Together, these results suggest synaptic histamine may adaptively shift value updating toward a more stable (or, less reactive) state in aversive contexts.

### Comparative H3R-weighting, cerebral blood perfusion, blinding integrity and subjective affect

Additional control analyses were performed to examine H_3_R specificity and potential confounds in the primary analyses. Applying H_3_R-weighting to task-based fMRI clusters (pitolisant > placebo) increased the magnitude of group-level effects (mean ± SE Δ*d* = +0.12 ± 0.05), with peak correspondence in the basal forebrain during new learning (Δ*d* = +0.34; Supplementary Fig. 15). Cerebral blood perfusion did not differ significantly between groups, either globally (grey and white matter) or regionally within task-specific networks (Supplementary Note 9; Supplementary Fig. 16).

Blinding integrity was assessed via participants’ estimation of group allocation and side-effect profiles. Group allocation estimates did not differ across groups at study completion (χ² = 0.00, *p* = 1.00, *N* = 58), with both groups predominantly guessing placebo (79%). Side-effect profiles did not differ significantly during acute administration (Supplementary Table 10).

As subjective affect and cognition (*e.g.*, motivation, arousal) can indirectly influence task behaviour ^76^, we assessed repeated measures of these domains. No significant group differences were observed over the testing period (Supplementary Fig. 17; Supplementary Tables 11 – 12).

## Discussion

Histamine has arguably remained the least understood of the canonical monoamines, despite being the first identified in the mammalian brain ^1–6^. The present findings help close this gap, revealing the causal role of histamine in the neurocomputational architecture of human learning. Together, these findings underscore histamine’s translational potential, especially in conditions characterised by cognitive impairment, where augmenting histaminergic signalling may counteract pathophysiological mechanisms ^77–80^.

Elevated synaptic histamine shifted temporal-hippocampal dynamics within a H_3_R-weighted network. This network-level separation was isolated to strengthened functional coupling between the hippocampus and mammillary zone – this zone encompasses both the histamine nucleus (TMN) and mammillary bodies, which share local circuitry and exhibit among the highest histaminergic fibre densities brain-wide ^13,81,82^. Notably, the mammillary bodies appear to function as a mediator of downstream histamine release in the hippocampus ^83–85^. The strength of hippocampal-mammillary zone coupling predicted both online encoding-related activity within the hippocampus, and post-encoding signal persistence in the entorhinal cortex. Given synaptic histamine primes hippocampal CA3 sharp-wave activity ^86^, this raises the possibility that hippocampal-mammillary zone coupling may drive broader cortico-hippocampal dynamics in support of retrieval and consolidation.

Within the same temporal–hippocampal network where increased offline functional coupling was observed, elevated synaptic histamine increased neural recruitment during new learning, and sustained post-learning activity within the medial entorhinal cortex (MEC). MEC neurons exhibit persistent activity which directly modulates cortico-hippocampal activity ^87,88^, and disruption of MEC inputs during the post-learning period impairs memory formation ^89^. As synaptic histamine enhances theta-coupled spiking within the MEC ^31^, this points to a role for synaptic histamine in driving consolidation through sustained MEC signal persistence. Indeed, we observed that higher levels of MEC signal persistence predicted future retrieval computations, which in turn were asymmetrically shifted in those with elevated histamine levels (selectively enhancing evidence accumulation for previously encoded items, while reducing decision policy strength for distractors).

Learning systems interface with broader domains of cognition; accordingly, we probed the broader effects of elevated histamine on working memory and reinforcement learning. Under higher working-memory load, a shift in both computational strategy and dorsolateral prefrontal recruitment occurred in those with elevated synaptic histamine. Within the deep laminar layers of the prefrontal cortex where H_3_R is expressed ^90^, histamine release from TMN-projecting neurons regulates fast-spiking interneurons essential for aspects of working memory function ^91–93^. Potentially, H_3_R blockade via pitolisant shifted constitutive receptor activity within this network ^22,94^, contributing to the present findings. Accordingly, our findings complement but reverse the direction of prior human correlation work showing that greater DLPFC H_3_R density is associated with reduced *n*-back performance ^95^.

During reinforcement learning (RL), elevated histamine flattened learning rates for aversive outcomes. Preclinically, activation of H_3_R via optogenetics and TMN lesions produces the behavioural inverse of the present findings ^36–39^, while suppression of H_3_R via antagonism impairs aversive memory consolidation ^96^. These effects suggest that histamine influences aversive learning via circuits partly distinct from hippocampal-dependent memory, potentially through parallel engagement of cholinergic pathways ^9,96–98^. Accordingly, we found no relationship between RL parameters (*⍺*, *β*) and evidence accumulation efficiency *v* during memory retrieval (Supplementary Note 8), consistent with these effects arising from distinct computational mechanisms.

A common thread across the present findings is that synaptic histamine appears to bias neurocomputation toward stability and adaptive evidence accumulation, consistent with proposals that histamine supports novelty-linked arousal rather than undifferentiated global arousal ^23,99,100^. By selectively stabilising encoding-related activity and dampening reactive value updating, histamine may confer computational advantages in environments where relevant information must be retained and distractors filtered. However, as with other monoamines ^72,101–103^, histaminergic potentiation may incur broader costs, including anxiety and insomnia ^11,104,105^, counterbalancing its adaptive gating of novelty-linked arousal.

The principal mechanism of pitolisant is presumed to be upregulation of synaptic histamine via autoreceptor blockade ^42,46,106^, where it shows no intrinsic activity at H_3_R isoforms believed to function as heteroreceptors ^21,51^ (see Supplementary Note 10 for further discussion). Nevertheless, further work on the contribution of H_3_ heteroreceptors to human memory, and potential co-interaction with H_3_ autoreceptors ^107^, may offer important context to the present findings.

In conclusion, the present study highlights the broad mechanistic contribution of a long-neglected classical neurotransmitter system – histamine – to human behaviour. Specifically, we show that increased synaptic histamine shapes offline and online hippocampal dynamics, stabilising entorhinal activity post-learning and shifting future retrieval computations. More broadly, elevated histamine adapts neurocomputational strategy under high working-memory load while flattening learning rates for aversive outcomes. Given the co-occurrence of memory deficits in clinical populations characterised by histaminergic dysregulation, including depression and Alzheimer’s disease ^77–79^, investigating the translational potential of H_3_R blockade may be worthwhile.

## Methods

### Participants and design

Sixty healthy participants met the study eligibility and were randomised to receive a single dose of pitolisant or placebo (*N*=29:31, pitolisant:placebo). Study recruitment and data collection was undertaken between April 2023 and January 2024. To assess eligibility to participate, prospective participants underwent two screening sessions: an initial pre-screening (online) and subsequent in-person medical screening. Exclusion criteria included medical conditions which were contraindicated for pitolisant (*e.g.,* hepatic impairment), history of neurological or psychiatric health concerns, recent recreational drug use (3 month wash-out), neuroimaging contraindications, recurrent histaminergic drug use (*e.g.,* antihistamines for allergies), and current pregnancy or breastfeeding. All participants had a BMI between 18-31 and were fluent in English. To avoid functional (fMRI) confounds associated with handedness (*i.e.,* left- or right-handed) during cognitive memory performance ^26^, only right-handed individuals were included. Full study inclusion and exclusion criteria are described within Supplementary Table 1. For details of participant flow through recruitment and randomisation procedures, see the CONSORT flow chart (Supplementary Fig. 1). The final sample consisted of 58 healthy individuals (37 females; drug:placebo = 29:29; mean age = 28.17) and were well matched across baseline working memory (digit span), education (years and level of attainment), and other demographic factors (Supplementary Table 2).

Participants were randomised to receive a single high therapeutic dose of H_3_R inverse agonist pitolisant hydrochloride 36mg (18mg × 2 film-coated tablets) or placebo (lactose tablets), in a double-blind study design. As the experimental design centred on a multi-stage learning paradigm, a within-subject crossover was avoided as repeating the four-stage learning cycle may introduce meta-learning and period effects ^108,109^. Both pitolisant and placebo were encapsulated in a generic opaque capsule to conceal allocation from researchers and participants. Encapsulation and randomisation were undertaken by external researchers not engaged in data collection at the NIHR Oxford Health Biomedical Research Centre (Oxfordshire, United Kingdom).

Randomisation was undertaken using a novel variance minimisation algorithm which allows minimising potential group-level differences in general cognitive function at baseline ^110^. The algorithm balances group allocation while accounting for covariates – gender and baseline cognition score (calculated using the Wechsler Adult Intelligence Scale digit span aggregate ^111^) – using a computed sum of squared deviations.

The study was assessed and approved by the Central University Ethics Committee at the University of Oxford (MSD-IREC reference code: R83940/RE002). All participants provided informed consent prior to study participation. The study and its protocol were pre-registered on the National Institute of Health Clinical Trials Database (NCT05849675). All pre-registered primary outcomes are presented in this paper. Screening and study visits were conducted within the Department of Psychiatry and Oxford Centre for Human Brain Activity, University of Oxford.

### Procedure

Participants undertook an online pre-screening procedure to assess basic eligibility (*i.e.,* BMI, medical status and handedness) before being invited to an in-person medical screening. During the in-person screening, participants were administered the Structured Clinical Interview for DSM-V to screen for current or past psychiatric illness. Medical history, medications, and MRI contraindications were assessed in interview. Blood pressure and heart rate was taken, and urine samples were acquired to assess current pregnancy and/or drug use. During this visit, participants undertook the Wechsler’s Digit Span test (forward, backward and ordered) ^111^, an index of general intelligence/cognition (Gignac & Weiss, 2015; Hale et al., 2002; Jones & Macken, 2015; M. Martin, 1978), which allowed balancing general cognitive functioning across allocation groups (via variance minimisation randomisation).

Eligible participants who passed the screening process were invited to the study visit, where the main study outcome data was collected. At the start of the study visit, participants completed baseline questionnaire battery of affect (for further details on questionnaire measures, see Supplementary Methods). Following this, participants self-administered their assigned intervention (pitolisant or placebo) in the presence of a researcher. Next, there was a three-hour period to allow drug metabolism before commencing neuroimaging and behavioural experiments. After three hours, pitolisant reaches peak serum concentration and high occupancy of brain-wide H_3_R within the high therapeutic dose range used within the present study ^50,116^. After three hours, participants undertook neuroimaging, behavioural testing and questionnaire procedures at the Oxford Centre for Human Brain Activity. At the end of the study visit, participants were asked to estimate their allocation before study debriefing.

### fMRI memory and learning task paradigms

The multi-stage memory paradigm (Fig. 1A; adapted from ^117^) is composed of four distinct stages. First, participants engage in the encoding of visual stimuli (images of landscapes or animals) through repetitive encoding. To support encoding, participants categorised images by content via button press (‘A’ for animal, ‘L’ for landscape). In the second stage, resting-state fMRI (rsfMRI) was undertaken over a 10.4 minute interval to capture network dynamics during the post-learning stage. The third stage involves a second encoding task similar to the first; however, it incorporates both previously learned images (*n* = 8) and novel images (*n* = 48) within an fMRI scanning session. This design enables direct comparison of neural responses associated with the encoding of new information versus familiar content (i.e., novel > familiar contrasts); task sequences were optimised to probe the hippocampal subfields (dentate gyrus and cornu ammonis [CA1-3]) given their importance in memory formation ^117,118^. Finally, the fourth stage comprises a memory recognition task in which participants are asked to determine whether each presented image was encountered earlier in the experiment (from either the first or third stage; *n* = 56) or represents a novel distractor (*n* = 27). During this task, participants indicate if images were previously seen via button response (‘A’ for previously seen, ‘L’ for unseen distractors). For the memory recognition task, behavioural task outcomes (accuracy and response time) were used to undertake drift diffusion modelling of evidence accumulation (see Supplementary Methods for further details).

The complex working memory task (verbal *n*-back; Fig. 4A) captures neural activity during working memory processing under increasing task complexity, inducing activation changes within the DLPFC, hippocampus and dopaminergic nuclei ^119,120^. During the task, participants must identify if a letter corresponds to a previously presented letter according to a rule. Specifically, if the letter occurred one trial ago (1-back), two trials ago (2-back), or three trials ago (3-back). In addition, there was an intermixed sensorimotor control task which served as a performance baseline (0-back). Letters appeared interleaved in sets of ten trials per block with four blocks per task condition (16 task blocks and 17 rest blocks total). To deter phonological and visual task strategy, occlusive consonants served as task stimuli in both upper and lower case form. The behavioural task outcomes were adjusted overall accuracy (correct hits – false alarms) and response time. Additionally, behavioural data for the task was fit to a computational model of evidence accumulation, a drift diffusion model, which was validated here through model selection, parameter recovery, and posterior predictive checks (see Supplementary Methods for further details). The winning model contained the following parameters: drift rate (+***v***/−***v***), non-decision time (***T***_***er***_) and decision policy (***a***). This task, associated preprocessing scripts, and computational modelling scripts are shared via an open-source repository: https://github.com/mjcolwell/n-back_oxford.

The Probabilistic Instrumental Learning Task (Version 220916; adapted from ^71,73^) assesses reward and loss-based instrumental learning using a fixed-probability outcome structure. Participants chose between two symbols in alternating pairs: one pair associated with high-probability reward trials (outcomes: +£0.20 or no change), and the other with high-probability loss trials (outcomes: –£0.20 or no change). Each symbol within a pair was associated with a fixed outcome probability (70% vs. 30%), and trial outcomes were presented immediately following the choice. Participants were informed prior to the task that their cumulative earnings would be awarded upon study completion and were instructed to maximise monetary gain. Primary behavioural outcomes included the proportion of optimal choices (i.e., selecting the option with the higher probability of gain or loss avoidance) and response time (ms). In addition, computational Q-learning models were fit to choice behaviour, allowing access to parameters (learning rate *⍺*; inverse decision temperature *β*) which formalise decision learning and computations throughout the task (for further details, see Supplementary Methods).

### Behavioural and demographic data analysis

Behavioural task and demographic data pre-processing and statistical analyses were carried out using R Software (version 4.3.1) ^121^. The effect of group allocation (pitolisant vs. placebo) on behavioural task outcomes was undertaken via two-way mixed-effect ANOVA modelling, with group and task condition serving as fixed factors and participant as a random factor; the variance minimisation procedure allows for ANOVA analysis without inclusion of balancing covariates ^110^. Planned comparisons were undertaken via two-tailed estimated marginal means analysis (EMM), where estimates are reported with associated standard means error. Family-wise error (FWE) was corrected for using the Bonferroni-Holm procedure. Effect size metrics are reported for ANOVA (partial eta squared: η_p_^2^) and EMM (Cohen’s d, *d*) alongside the associated two-tailed 95% confidence intervals (effect size calculations are described in the Supplementary Methods). Variables with skewness of ≥ ±1 or kurtosis ≥ ±5 were log-transformed, and underwent confidence interval bootstrapping for EMMs tests. In addition to normality checks, models were verified for covariate independence and homogeneity of regression slopes by assessing covariate interactions across factors. Homogeneity of demographic covariates across groups was assessed using chi-squared (X^2^) independence tests (binary or categorical data) or independent two-tailed t-tests (discrete or continuous data). Participant group estimation results were analysed via X^2^ independence tests. Inferential analyses were carried out at an alpha level of 0.05. A sample size calculation determined a required sample size of *N*=52 was needed to undertake two-tailed between-groups inferential statistics (Power [1 - β error probability]: 80%).

### MRI data acquisition, preprocessing and analysis

Multi-modality MRI data was acquired using a 3-Tesla Prisma Siemens scanner utilising a 32-channel head matrix coil at the Oxford Centre for Human Brain Activity (University of Oxford). The scan started with T1-weighted anatomical image acquisition across 192 slices (TR = 1900ms, TE = 3.97ms, field of view = 192mm, flip angle = 8°, voxel dimensions = 1mm^3^, acquisition time = 5 mins, 31 sec). Functional image sequences were T2-weighted echoplanar images and were parameterised to optimise for image fidelity in regions of interest (ROIs). The first task, the novel memory encoding task, was acquired over one run (TR = 800ms, TE = 30ms, flip angle = 52°, slice thickness = 2mm, multiband accelerator factor 6, resolution = 2.4mm^3^ isotropic voxel size, acquisition time = 6 mins, 48 sec). The second task, the complex working memory task, was also acquired over one run (TR = 1500ms, TE = 25ms, flip angle = 70°, slice thickness = 2mm, multiband accelerator factor 3, PAT GRAPPA acceleration factor 2, resolution = 2mm^3^ isotropic voxel size, acquisition time = 15 mins, 9 sec). To assist in distortion correction of functional images, fieldmaps were acquired before task runs (TE1 4.92ms, TE2 7.38ms; TR = 590ms, flip angle = 46°).

Structural and functional data were converted to BIDS specification using *HeuDiConv* (v 1.0.1)(https://github.com/nipy/heudiconv) and converted to NIfTI-1 using *dcm2niix* (v1.0.20220720) (X. Li et al., 2016). Image quality for structural and functional data was assessed using *MRIQC* (v23.1.0) ^123^. Structural images were defaced using *pydeface* (v2.0.2) (https://github.com/poldracklab/pydeface), bias-field corrected using *FSL* (version 6.00) ^124^, deep-learning based skull-stripping via *synthstrip* (v1.3) ^125^. All functional images were registered to T1-weighted images using FSL’s *FLIRT/FNIRT* ^126^ and normalised to Montreal Neurological Institute (MNI) space.

Functional resting state data underwent first-level preprocessing via FSL’s *MELODIC* tool: high pass temporal filtering occurred at 99s, motion correction was undertaken via MCFLIRT ^127^, field-map correction, and automatic dimensionality estimation of components. These pre-processed data were further denoised with a supervised learning approach via ICA-FIX ^128,129^. To account for the specific acquisition sequence used, the ICA-FIX model was self-trained on signal/noise discrimination via hand-labelling of 16 participants from this dataset; leave-one-out (LOO) cross-validation produced a 3 × *TPR* + *TNR* of 99.9% at a threshold of 30, which was the threshold used for denoising the full dataset. Following denoising, the data were smoothed using a Gaussian kernel of 2.5mm and normalised to MNI space.

To investigate how increased synaptic histamine influences offline network dynamics, *a priori* ROIs for network nodes were defined. ROIs were restricted to regions with strong evidence for both (*i*) involvement in memory processing (and thus also used in the memory encoding task), and (*ii*) sensitivity to histamine H_3_R modulation, based on prior animal and human work ^13,31,32,39,53–59^. This approach reduced dimensionality and minimised the risk of spurious network effects. Additional temporal-limbic regions were excluded where no evidence for H_3_R involvement was available. An ROI of the bilateral hippocampus was derived from the Jülich histological probabilistic atlas ^130^, containing a combined bilateral mask of the dentate gyrus and cornu ammonis (CA) (set to a > 0.50 probability threshold). An *a priori* ROI of the mammillary zone, which contains with the tuberomammillary nucleus (origin of histamine neurons) as indicated past work ^61,62^, was derived from the Jülich atlas (> 0.25 probability threshold) ^131^. Additionally, an *a priori* ROI of the basal forebrain was also derived from the Jülich atlas ^130,132^. Further ROIs of the bilateral entorhinal cortex (set to a > 0.50 probability threshold), also derived from the Jülich atlas ^130,132^, and bilateral perirhinal cortex masks were created using 5mm radius spheres with reference to previous work on this region (MNI coordinates: left mask centre X,Y,Z [-33, -4, -32]; left mask centre [33, -7, -29]) ^133^. To correct for multiple comparisons, *a priori* ROIs were bilateralised/contained all divisions. During the post-learning resting stage, time series signals were extracted from these ROIs; extracted signals were multiplicatively weighted using a rank-based inverse normalised PET image of regional H_3_R density from neuromaps ^60^. These images were originally supplied by Gallezot *et al*. ^24^ from a sample of participants with a mean age of 35 ± 10 (*N*=9; all male), within 6 years of the mean age range within the current study.

Covariance network matrices of H_3_R-weighted regions of interest (ROIs, or nodes) were generated using *fslnets* (http://fsl.fmrib.ox.ac.uk/fsl/fslwiki/FSLNets). Multivariate analysis of these covariance matrices was performed using linear discriminant analysis, applied to regularized partial correlation matrices derived from all participants’ data (from *scikit-learn* ^134^). This machine learning algorithm employs LOO cross-validation to train and test classifiers, enabling the discrimination of network matrices between different allocation groups. These analyses were also performed on the non-weighted dataset and are reported in Supplementary Note 1.

Task-related functional data were preprocessed and analysed (first-level) using FSL’s *FEAT* tool: motion correction was undertaken using *MCFLIRT* ^127^, with spatial smoothing set to a Gaussian kernel of 5mm (full-width-half maximum), high-pass temporal filtering (fitting a straight line using Gaussian-weighted least squares; memory encoding task σ = 45.0s; working memory task σ = 49.5s), field-map correction, normalisation of the 4D dataset grand-mean intensity using a single multiplicative factor, and distortion correction via B0 unwarping using phase and magnitude images. For task-related functional images, motion outlier confound matrices were created using *FSL*. During registration, functional images were aligned to a single-band functional reference image with increased contrast acquired before the task acquisition sequence. Functional image datasets with high motion (maximum absolute displacement of ≥ 1.5mm) or technical issues during the scan (*e.g.,* scanner failure during time-dependent sequence) were excluded from the dataset (*N*=6 for all functional sequences). H_3_R-weighting was not applied to task-based fMRI analyses, as the permutation-based GLM pipeline (FSL’s randomise) does not permit integration of external PET-derived weights. We therefore prioritised analysis of unweighted task-related neural signal, before examining the histaminergic specificity of associations via post hoc application of receptor-weighting to extracted parameter estimates (see Supplementary Methods).

For task-related functional data, higher level analysis was undertaken using non-parametric permutation inferential analysis via FSL’s randomise ^135^, utilising the threshold-free cluster enhancement (TFCE) method set to *p* < 0.05. Significant clusters identified through TFCE were summarized using descriptive statistics of the peak voxel within each cluster (i.e., location, t-statistic and p-value). Motion outlier confound matrices and motion parameters were included in higher-level analysis. The main effect of task was analysed at the whole brain level, while the group-level analyses (pitolisant vs. placebo) were restricted to *a priori* ROIs which overlap with histological maps of histaminergic innervation (small volume correction [SVC]) ^13,32,53,54,136^. Follow-up exploratory analyses for both tasks undertaken using a whole-brain mask. For the novel memory encoding task, neural activity during new learning was contrasted with previously learned information (novel > familiar contrast). In addition, BOLD signal from each rest period (12s) was decomposed using finite impulse response across 15 temporal bins (one bin per 0.8-s TR). These bins were concatenated within a linear model which captured signal decay which contrasted rests following novel > familiar encoding (*t*_1_ novel [EV -7.5] → *t*_15_ novel [EV +6.5]; *t*_1_familiar [EV +6.5] → *t*_15_familiar [EV -7.5]). For the complex working memory task, increased activity as a function of increased task complexity/load was linearly modelled (0-back [EV -1.5], 1-back [EV -0.5], 2-back [EV +0.5], and 3-back [EV +1.5]). Alongside this, contrasts across each level of task complexity were undertaken: 1 > 0-back, 2 > 0-back, and 3 > 0-back. During the memory encoding and complex working memory tasks, the same *a priori* ROIs from the network analysis were reapplied: bilateral hippocampus, basal forebrain, mammillary zone, entorhinal and perirhinal cortices. Each ROI was bilateralised/contained all divisions. For the complex working memory task, ROIs were created over the bilateral medial DLPFC (BA46 and dorsal transition area BA46/9) and bilateral rostral DLPFC (BA9), per previous subdivisions of this region and task relevance ^137,138^, using cortical maps from the Human Connectome Project atlas ^139^. Given functional differences across hemispheric divisions of the DLPFC in working memory function ^137,140–142^, each division was treated as distinct. In addition, given the role of the dopaminergic substantia nigra in n-back performance, alongside its histaminergic innervation ^120,136^, an ROI mask derived from previous work was selected (set to a > 0.25 probability threshold) ^143,144^. All ROIs were resampled to 2mm^3^ MNI space and were previously investigated at 3T fMRI ^120,133,137,144–148^. Significant clusters for each ROI are reported alongside their corresponding peak MNI coordinates. Parameter estimates from significant clusters were used to assess their relationship with experimental variables such as connectivity values (see Supplementary Methods for details). The voxel-wise statistical maps resulting from permutation testing were processed and visualised using *Nilearn* (v 0.10.4) (https://github.com/nilearn/nilearn) and *Nibabel* (v 5.2.1) (https://github.com/nipy/nibabel).

Arterial Spin Labelling (ASL) was undertaken to explore the potential influence of pitolisant on cerebral blood flow. Details on the ASL sequence parameterisation, preprocessing and analysis are provided in the Supplementary Methods.

## Data Availability

Raw and modelled datasets for this study have been deposited on Github: https://github.com/mjcolwell/Histamine_Learning_Data_and_Code

## Code availability

The code used to undertake preprocessing, network analysis, computational modelling and inferential modelling are available on Github: https://github.com/mjcolwell/Histamine_Learning_Data_and_Code

## Acknowledgments

This work was funded by the NIHR Oxford Health Biomedical Research Centre and the NIHR Oxford Cognitive Health Clinical Research Facility. The views expressed are those of the authors and not necessarily those of the NHS, the NIHR or the Department of Health.

We thank the radiography team at the Oxford Centre for Human Brain Activity, University of Oxford (Nicky Aikin, David Parker, Juliet Semple, Michael Sanders, Jon Campbell and Camille Lasbareilles). We thank Dr Nicola Filippini for providing task materials. We thank Dr Sebastian Rieger for technical assistance throughout the study period. We thank Anutra Guru, Esther Teo, and Dr Wendy Howard for supporting the variance minimisation procedure. We thank Dr Jan Grohn for suggestions for data analysis. We thank Dr Thomas Okell for providing ASL sequence parameterisation.

## Author Contributions Statement

M.C. conceived and designed the study, with additional input from senior authors (P.C.; S.M.; C.H.). All analyses (neuroimaging, computational, and behavioural) were conducted by M.C. M.C. produced all figures and drafted the manuscript. Data collection was undertaken by M.C. and F.V.U., with technical input from M.B. and M.M. All authors contributed to revisions and approved the final manuscript.

## Competing Interests statement

C.H. has received consultancy fees from P1vital Ltd., Janssen Pharmaceuticals, UCB, Compass Pathways, and Lundbeck. She is a co-director of TnC Psychiatry and Neuroscience. S.M. has received consultancy fees from Zogenix, Sumitomo Dainippon Pharma, P1vital Ltd. and Johnson & Johnson Pharmaceuticals. C.H. and S.M. recently held grant income from Zogenix, UCB Pharma and Janssen Pharmaceuticals. C.H., S.M. and P.C. recently held grant income from a collaborative research project with Pfizer. M.B. reports grants from the Wellcome Trust, Medical Research Council, Office for Life Science and National Institute for Health and Care Research (NIHR); consulting fees from Janssen, CHDR, Engrail Therapeutics, Boehringer, P1vital, Alto Neuroscience, and Empyrean Therapeutics; and was previously employed by P1vital. The other authors report no conflicts of interest.

