## Supplementary Materials for "Synaptic histamine shapes the neurocomputational dynamics of human learning"

*Colwell et al. N.D.*

#### Supplementary Methods

##### Questionnaire Battery, Procedure, and Analysis

A battery of self-report questionnaires was administered at baseline (i.e., the start of the study visit) and again at the end of the visit to assess self-rated state anxiety, anhedonia, affect, and side effects. These included the Spielberger State Anxiety Inventory (STAI-S) [1], the Dimensional Anhedonia Rating Scale (DARS) [2], the Positive and Negative Affect Schedule (PANAS) [3], a Visual Analogue Scale (VAS), and a side effects profile questionnaire. As side effects were recorded only during the study visit, items relating to insomnia and sexual side effects were excluded from analysis.

In addition to these established measures, a novel instrument – the State-Subjective Cognition Scale (SSCS) – was employed to assess subjective cognitive states. The SSCS was developed specifically for this study to address limitations of existing measures of subjective cognition (Van Umm, Colwell, & Murphy, in submission). It comprises fifteen items rated on a visual analogue scale (0–100), grouped into five psychometrically validated subscales: motivation, mood, executive function, selective attention, and memory. The SSCS was administered at four time points during the study visit: at baseline ($T_1$), prior to MRI scanning (2.5 hours post-dose; $T_2$), immediately following MRI scanning (4 hours post-dose; $T_3$), and at the conclusion of the visit ($T_4$).

For the main questionnaire battery (excluding the SSCS), post-dose questionnaire scores were analysed using baseline-adjusted mixed-effects ANCOVA models. In these models, post-intervention scores served as the outcome variable, with baseline (pre-intervention) scores included as a covariate, and participant was included as a random effect. Given the added temporal dimension of the SCSS ($T_{1\text{-}4}$), these data were analysed using linear mixed-effects models with restricted maximum likelihood estimation, incorporating time as a fixed effect. As with task data, planned pairwise comparisons for questionnaire data were conducted using estimated marginal means with two-tailed tests, and multiple comparisons were corrected using the Bonferroni-Holm procedure. All inferential tests were evaluated at an alpha level of 0.05.

##### Memory Recognition Task Analysis – Drift Diffusion Modelling

The memory recognition task, conducted during the final phase of the multistage fMRI memory paradigm, assessed participants' memory for previously encoded visual items among novel distractors. Task behaviour was fit to a computational model of evidence accumulation, a Bayesian drift diffusion model (DDM). The DDM provides access to latent cognitive processes which occur during evidence accumulation, providing potential explanations for patterns observed within task behaviour [4]. Our DDM approach utilised the publicly available *HDDM* [5] package available in Python, which allows undertaking Bayesian Monte Carlo Markov Chain (MCMC) sampling (via *PyMC* [6]) of all model parameters per participant. Each model was calculated with four Markov chains with 4,000 samples per chain, of which the first 2,000 were discarded as burn-in. Models were assessed for convergence via *ArviZ* [7] using the Gelman-Rubin diagnostic (rank-normalized, split-$\hat{R}$); the threshold for acceptable convergence was set to $\hat{R} \leq 1.01$. Model selection was guided by the deviance information criterion (DIC), which balances model fit and complexity [8]. The initial candidate model was the simplest possible DDM, consisting of three parameters: drift rate parameter ($v$), decision policy ($a$), trial variability, and non-decision time ($T_{er}$). Model selection occurred through iteratively increasing complexity through a stepwise approach (Supplementary Table 5). Allowing parameters to vary according to stimulus type ('previously encoded' or 'distractor'), and the inclusion of inter-trial variability parameters resulted in better model fit. The stimulus-dependent estimation of parameters aligns with previous implementations of the DDM for recognition memory [9]. The winning model consisted of four stimulus-dependent parameters, $v_s$, $a_s$, $T_{er,s}$ and $z_s$, with two fixed inter-trial variability parameters: $st$ and $sz$. This model can be conceptualised as following:

$$\mathrm{RT}(x) \approx \frac{a_s{}^{eff}(\, z'_s)}{v_s} + T_{er,s} \; + \; \mathcal{N}(0, st^2)$$

where the trial-wise starting point $z'_s$ varies according to:

$$z'_s \sim \text{Uniform}\,(z_s - \frac{sz}{2}, z_s + \frac{sz}{2})$$

Where the differential varies according to if the lower boundary is hit $a_s{}^{eff} = \; a_s \cdot \; z'_s$ , or if the upper boundary is hit $a_s \cdot (\, 1 - \; z'_s)$. $v_s$ was sampled in unconstrained Euclidean space, while $a_s$ , $z_s$ , $sz$ and $st$ were sampled in space and transformed to positive reals. $T_{er,s}$ was sampled within a space bounded by the observed RT range. ). Model parameters were estimated for individual participants without access to group-level posteriors to allow frequentist statistical analyses. Posterior predictive checks indicated a close match between synthetic data retrieved from the model fitting process and observed behaviour (Supplementary Fig. 5). Parameter recovery for stimulus-dependent and fixed parameters was performed using *pyMC* [6] MCMC, using true parameter estimates across 200,000 trials. All parameter estimates were checked for recoverability (Supplementary Fig. 6), while the winning model demonstrated adequate convergence during fitting ($\hat{R} \leq 1.01$). Before inferential analyses, decision policy $a_s$ was log transformed, $z_s$ was inverse logit transformed, while $T_{er}$ was arcsine transformed. All parameters were analysed using mixed-effects ANOVA and EMM modelling approaches.

#### *N*-Back outcome analyses and drift diffusion modelling

Due to excess skewness (-1.67 [threshold ≥ ±1]) and kurtosis values (6.25 [threshold ≥ ±5]), raw accuracy (%) values on the *n*-back were log-transformed, and EMM tests included bootstrapped confidence intervals. Raw accuracy (%) values are retained within Fig. 4D (main text) for interpretability.

A Bayesian drift diffusion model (DDM) was validated for the verbal *n*-back task as part of the present study. As with the memory recognition task, the DDM was fit via HDDM [5] with each model being calculation across four Markov chains with 4,000 samples per chain, of which the first 2,000 were discarded as burn-in. In addition, model convergence was assessed using a split-$\hat{R}$ criterion of ≤ 1.01. The initial candidate model was selected based on previous work with the DDM which included a drift rate parameter ($v$), an initial starting bias parameter ($z$), decision policy ($a$), trial variability, and non-decision time ($T_{er}$) [10]. Due to the high trial type imbalance (20-30% cued targets per block), we considered models with separate drift rates ($+v/-v$). In our model comparison, the best-fitting model included split drift rate but excluded initial choice bias, as it yielded the lowest Deviance Information Criterion (Supplementary Table 8).

Drift criterion for split drift rate was not included as there was no theoretical basis for its influence in this paradigm [10]. While more complicated models were considered, high parameterisation has been reported to reduce detection of group-level effects from DDM parameters [11]. The winning candidate model ($+v/-v,\ a,\ T_{er},\ st,\ sv$) can be conceptualised as following:

$$\mathrm{RT}(x) \approx \frac{a}{v + \mathcal{N}(0,\ sv^2)} + T_{er} + \mathcal{N}(0, st^2)$$

As with the memory recognition task, all models parameters ($v, a,\ st$ and $sv$) were sampled in Euclidean space, except $T_{er}$ which was sampled in a space bounded within the observed RT values. Model parameters were estimated at the individual level without incorporating group-level posteriors, enabling subsequent frequentist statistical analyses. Posterior predictive checks showed a close match between observed task and synthetic data derived from the model fitting process for participant data (Main text, Fig. 4C). Parameter recovery was undertaken using synthetic data simulated via *pyMC* MCMC using true parameter estimates for each model parameter across 200,000 trials. These simulated data were fitted using the winning DDM, and recovered parameter values were compared against their true values. All model parameters were deemed recoverable from the fitting process (Supplementary Fig. 8A). The winning model converged well during the fitting process ($\hat{R}$ ≤ 1.01). For inferential analyses, decision policy $a$ was log transformed, while $T_{er}$ was arcsine transformed. All parameters were analysed using mixed-effects ANOVA and estimated marginal means (EMM) modelling approaches.

Using simulated task data from the posterior predictive checks, we performed post-hoc analyses to assess if high and low $T_{er}$ (split using within-subjects median-split) influence posterior accuracy. We ran a mixed-effects ANOVA model with $T_{er}$ bins as a between-subjects factor and posterior accuracy as the outcome variable and estimated marginal means planned comparisons.

#### Probabilistic Instrumental Learning Task Analysis and Reinforcement Learning Modelling

For the analysis of behavioural performance, the final half of trials (10 per condition) were analysed, as this reflects the plateau phase of learning performance [12,13]. One participant was excluded due to data quality issues. Exploratory analyses including all trials is reported in Supplementary Note 9, alongside analyses comparing overall monetary gain across allocation groups.

Behavioural data were fit using a computational reinforcement learning model (Q-learning), consistent with prior implementations [12,14,15]. Model selection details are provided in [14], with parameter recovery checks reported in [12]. Posterior predictive checks were conducted by comparing observed temporal learning patterns with those simulated from the fitted models (see Supplementary Fig. 14D; Main text, Fig. 5B). The value of a selected stimulus was updated on each trial, with the value of the unselected stimulus updated reciprocally. The learning algorithm is described in the following equation:

$$Q_{t+1(s)} = Q_{t(s)} + \alpha_j \left(R_t - Q_{t(s')}\right)$$

The Q value, or belief expectation, $Q_{t(s)}$, describes the value associated with a symbol ($s$) at a given trial ($t$). $R_t$ refers to the type of outcome observed, which can be positive (win during high-probability win [1], or avoiding loss during high-probability loss [0]) or negative (no change during high-probability win [0], or loss during high-probability loss [-1]). The learning rate parameter $\alpha_j$ , which determines the slope of the learning curve, is set for each trial type $j$ (win/loss trials) and was sampled in logit space. In line with previous studies [14], $Q_{0(s)}$ was initialised at 0.5. Each trial, the unselected option, described as $Q_{t(s')}$, was updated reciprocally based on the counterfactual outcome from the selected symbol. The Q value assigned to the first two paired stimuli were used to provide an estimate of choice probability through a softmax function:

$$P_{t(s)} = \frac{1}{1 + exp^{-\beta_j\left(Q_{t(s)} - Q_{t(s')}\right)}}$$

The probability ($P_{t(s)}$) of choosing a given the shape $s$ is controlled by the inverse decision temperature parameter $\beta$ (sampled in log space). As with the learning rate parameter, $\beta$ is set for each trial type $j$. Modelling win and loss trials separately was required to observe valence-specific effects and is in line with previous implementations of the model [12]. Each parameter was estimated within discrete space (110 × 100 grid) by determining the posterior probability of each trial type, then by calculating the expected value for each marginal likelihood parameter. The $\alpha$ parameter was log transformed before inferential analyses using mixed-effects ANOVA and planned comparisons EMM tests. Given the high covariance between model parameters due to within-subjects trade-off, mixed-effects ANOVA models were adjusted to include the reciprocal parameters as random effect terms: `(parameter (α/β) ~ group * condition (loss/win) + (1 + parameter (α/β) || Participant))`. Independent random intercepts were specified to account for condition-level parameter trade-offs, as computational models were fit separately for each condition within participants.

Beyond the computational model reported, alternative models were considered; these include models where outcome sensitivity and value decay are parameterised. However, these parameters are mathematically redundant [16], therefore a theoretical prior to justify their inclusion. Given the lack of previous work on histamine and reinforcement learning, it was decided to use the legacy Q-learning parameter $\beta$ [15].

#### Global and Regional Blood Perfusion Analyses

Following rsfMRI and task-based fMRI sequences, an arterial spin labelling sequence occurred to estimate global and regional blood flow across allocation groups. Participant structural data was preprocessed used FSL's anatomical processing script (*FSL_Anat*; https://fsl.fmrib.ox.ac.uk/fsl/fslwiki/fsl_anat) which entailed bias-field correction, brain extraction and registration to standard MNI152 space via FMRIB's *FNIRT* tool). ASL perfusion maps were distortion-corrected and motion-corrected within units of mL/100mg/min via FSL's *BASIL* (Bayesian Inference for Arterial Spin Labelling tool, https://fsl.fmrib.ox.ac.uk/fsl/fslwiki/BASIL) [17,18]. BASIL performed voxelwise calibration, label-control subtraction, and inferrential estimation of voxel-wise perfusion to generate absolute perfusion maps, controlling for the partial volume effects observed at the participant level. These perfusion images were then registered to standard space using the non-linear warp generated via *FSL_Anat* via *FNIRT*. The transformed and calibrated perfusion maps were analysed using global CSF values output by BASIL for grey and white matter and analysed via between-groups ANOVA modelling. For regional CSF analysis, voxelwise glm permutation testing (5000 permutations, with TFCE and FWE correction for cluster significance) was undertaken within whole-brain and task-network maps via FSL's *randomise*. For inferrential analyses of calibrated perfusion maps, there were missing data for one participant (due to scanner sequence issues) and exclusions due to technical issues during the scan (*N*=5). Global estimates of blood flow analyses included grey and white matter masks. Regional estimates of blood flow were undertaken within a mask of *a priori* region of interests for the fMRI sequences (memory consolidation/encoding and complex working memory).

#### fMRI Resting State and Task-Based Correlational Analyses

Linear regression analyses (including two-tailed Pearson's product-moment tests) were conducted to examine the relationship between group-level effects observed across outcome variables (including fMRI TFCE-corrected cluster β values and computational parameters) within the primary analysis. The relationship between Mammillary zone–hippocampal connectivity (predictor) and both hippocampal encoding cluster activity (outcome) and entorhinal memory trace cluster (outcome) was assessed (see main text, Fig. 2C – G, I). In addition, the relationship between trace decay (predictor) and computational decision parameters during recall (drift rate $v_s$ and decision policy $a_s$) was assessed. For the *n*-back task, the relationship between drift rate $v$ (outcome) was assessed with activity within the left DLPFC cluster as task complexity increased (predictor). While a significant group effect was observed for non-decision time $T_{er}$ (see main Results), there was no theoretical basis for its examining its relationship with DLPFC activity given activity within this region has been causally linked to $v$ across multiple studies, but not $T_{er}$ [19,20]. Where appropriate, correlational analyses were corrected for multiple comparisons using Holm–Bonferroni adjustment. For each linear regression analysis, follow-up models were fitted separately within each allocation group using ordinary least squares. This allowed estimation of the predictor–outcome relationship independently within groups, without assuming a shared effect across conditions. Regression coefficients from these within-group models were then aggregated to characterize the posterior distribution of effects across groups.

For all correlational analyses involving fMRI signal, the signal was weighted based on the spatial density of regional $H_3R$ (rank-based inverse normalized). A comparison between weighted and non-weighted group-level effect sizes is reported in Supplementary Fig. 15. In addition, a comparison between weighted and non-weighted correlational analyses is reported in Supplementary Table 9.

#### Effect Size Calculation

Effect sizes were derived for both mixed-effects ANOVA models and estimated marginal means (EMM) comparisons. For ANOVA models, partial eta squared ($\eta p^2$) was computed using the *eta_squared* function from the *effectsize* R package, applied to the fitted mixed-effects model. For EMM pairwise comparisons, Cohen's *d* was estimated using the *eff_size* function from the *emmeans* R package, based on follow-up outcome values.

#### Behavioural Data Software – Collection and Analysis

For data collection, the following software packages were used: *PsychoPy* 2021.1.4; *MATLAB* 2021a; *Qualtrics Surveys*; *Anaconda* 2021.4; *Presentation* (Neurobs) 23.0; *Psychtoolbox* v3. For data analysis, the following software packages were used: *Docker* 4.22.0; *MATLAB* (R2022a); *Python* (version 3.8.8); R (4.3.1). The following R packages were utilised: *moments* (0.14.1); *effectsize* (0.8.6); *ggbeeswarm* (0.7.2); *gtools* (3.9.4); *readxl* (1.4.2); lme4 (1.1-33); *sdamr* (0.2.0); *plotrix* (3.8-2); *ggridges* (0.5.4); *tidyverse* (2.0.0); *knitr* (1.42); car (3.1-2); *stringr* (1.5.0); *ggplot2* (3.4.2); *ggsignif* (0.6.4); *ggpubr* (0.6.0); *viridis* (0.6.4); *ggdist* (3.3.0); *ggstance* (0.3.6); *RColorBrewer* (1.1-3); *ggrepel* (0.9.3); *emmeans* (1.8.5); *ggpp* (0.5.4); *lmerTest* (3.1-3); *cowplot* (1.1.1); *openxlsx*

(4.2.5.2); *ez* (4.4-0); *gghalves* (0.1.4); *ggsignif* (0.6.4); *data.table* (1.14.8); *ggplot2* (3.4.2); *psycho* (0.6.1); *gtools* (3.9.4); *ggExtra* (0.10.1); . All required Python dependencies are included within the Docker image: hcp4715/hddm:0.8. No additional MATLAB packages required.

#### Supplementary Note 1

Following iterative encoding of visual stimuli in phase one of the multi-stage fMRI task, a short test was undertaken to ensure a high accuracy criterion was achieved (≥85% accuracy). Participants discriminated between encoded items and novel distractors above criterion (mean = 98.98% ± 2.66). There was no significant difference in accuracy across allocation groups (Main effect of group [mixed effects ANOVA]: F[1,47] = 0.003, *p* = 0.957).

#### Supplementary Note 2

A network analysis without time series weighting using histamine 3 ($H_3R$) receptor maps was undertaken. Binary classification machine learning was able to differentiate between pitolisant and placebo with the same level of accuracy (88.46%). Connectivity between the mammillary zone and connectivity was higher in individuals who received the drug vs. placebo (*t*[51] = 2.79, *p* = 0.035, FWE-corrected, Cohen's *d* = 0.77).

#### Supplementary Note 3

During the formation of novel memories (novel > familiar contrast), there was increased activity across multiple brain regions in the pitolisant group, identified via *a priori* SVC permutation testing. These areas included the bilateral hippocampus (left hippocampus cluster: peak MNI coordinates = X,Y,Z [-22, -14, -26]; cluster size (voxels) = 89; *t*-max (51) = 4.651,*p* = 0.0054, TFCE-corrected; right hippocampus cluster 1: peak MNI coordinates = [22, -12, -26]; cluster size = 36; *t*-max (51) = 4.037, *p* = 0.0162, TFCE-corrected; right hippocampus cluster 2: peak MNI coordinates = [36, -26, -18]; cluster size = 6; *t*-max (51) = 3.724, *p* = 0.0396), basal forebrain (peak MNI coordinates = [6, 2, -16]; cluster size = 5; t-max (51) = 3.843, *p* = 0.0330, TFCE-corrected), bilateral perirhinal cortex (left perirhinal cortex cluster: peak MNI coordinates = [-34, -6, -34]; cluster size = 10; t-max (51) = 3.462, *p* = 0.0178, TFCE-corrected; right perirhinal cortex cluster 1: peak MNI coordinates = [36, 8, -34]; cluster size = 5; *t*-max (51) = 3.306, *p* = 0.0244, TFCE-corrected; right perirhinal cortex cluster 2: peak MNI coordinates = [38, -6, -28]; cluster size = 1; *t*-max (51) = 2.777, *p* = 0.0470, TFCE-corrected), and bilateral entorhinal cortex (right entorhinal cortex cluster 1: peak MNI coordinates = [22, -16, -28], cluster size = 53; *t*-max (51) = 4.256, *p* = 0.0016, TFCE-corrected; right entorhinal cortex cluster 2: peak MNI coordinates = [14, -6, -20], cluster size = 1; *t*-max (51) = 2.698, *p* = 0.0464, TFCE-corrected; left entorhinal cortex cluster: peak MNI coordinates = [-22, -16, -28], cluster size = 7; *t*-max (51) = 3.978, *p* = 0.0194, TFCE-corrected).

#### Supplementary Note 4

An analysis of labelling discrimination ('animal' or 'landscape') during the fMRI encoding task was undertaken to ensure visual stimuli were properly discriminated within the scanner. There was no significant main effect of allocation on labelling discrimination accuracy (F[1,50] = 1.72, $p$ = 0.195); in addition, there was no significant main effect of group on labelling speed (F[1,50] = 0.06, $p$ = 0.803) (Supplementary Fig. 2).

#### Supplementary Note 5

We examined whether the lateralisation of entorhinal trace persistence (main text, Fig. 2E – G) was mirrored by lateralised activity during encoding in significant clusters. There was a main effect of the left hippocampus (β = 0.54, $p$ = 0.004, $\eta_p^2$ = 0.17 [0.02, 0.36]; $r$ = 0.40) and left entorhinal cortex (β = 0.52, $p$ = 0.00921; $\eta_p^2$ = 0.14 [0.01, 0.34]; $r$ = 0.38) on signal trace persistence. However, there was no statistically significant main effect of the right hippocampus (β = 0.29, $p$ = 0.207) and right entorhinal cortex (β = 0.23, $p$ = 0.304).

#### Supplementary Note 6

Estimated marginal means (EMMs) for accuracy, drift rate, and non-decision time across n-back levels are reported here (as supplement to Fig. 4D, main text). Participants receiving pitolisant showed generalised increases in accuracy (0-back EMM ( ± SE) = 0.04 ± 0.03, $p$ = 0.115; 1-back EMM = 0.08 ± 0.03, $p$ = 0.0012, $d$ = 0.87 [0.30, 1.44]; 2-back EMM = 0.12 ± 0.03, $p$ = 0.0004, $d$ = 1.36 [0.84, 1.94]; 3-back EMM = 0.10 ± 0.03, $p$ = 0.0008, $d$ = 1.13 [0.60, 1.66]) and drift rate (0-back EMM = 0.42 ± 0.26, $p$ = 0.110; 1-back EMM = 0.63 ± 0.26, $p$ = 0.0163, $d$ = 0.89 [0.16, 1.62]; 2-back EMM = 0.48 ± 0.26, $p$ = 0.0632; 3-back EMM = 0.19 ± 0.26, $p$ = 0.452). As task complexity increased, non-decision time elevated in the pitolisant group (0-back EMM = 0.04 ± 0.03, $p$ = 0.1931; 1-back = 0.066 ± 0.032, $p$ = 0.04, $d$ = 0.78 [0.02, 1.54]; 2-back = 0.127 ± 0.032, $p$ = 0.0002, $d$ = 1.49 [0.72, 2.25]; 3-back = 0.11 ± 0.032, $p$ = 0.0009, $d$ = 1.30 [0.53, 2.06]). Splitting trials by high vs. low $T_{er}$ revealed decreased posterior accuracy under low load but improved accuracy under high load (0-back EMM = -0.03 ± 0.03, $p$ = 0.926; 1-back = -0.63 ± 1.75, $p$ = 0.004, $d$ = -0.63 [-1.38, -0.19]; 2-back = 0.02 ± 0.004, $p$ = 0.032, $d$ = 0.40 [0.04, 1.37]; 3-back = 0.03 ± 0.02, $p$ = 0.004, d = 0.69 [0.27, 1.52]).

#### Supplementary Note 7

We ran confirmatory analyses on the relationship between posterior *n*-back accuracy (mean %) and continuous $T_{er}$, using the following mixed effects model: `(`$T_{er}$ `~ accuracy*condition [reference = 0-back] + (1 | Participant))`. There was a significant interaction between accuracy and medium task difficulty (2-back; β = 1.25 [0.13, 2.38], $t$(176) = 2.20, $p$ = 0.0293) and the hardest task difficulty (3-back; β = 1.22 [0.11, 2.33], $t$(173) = 2.17, $p$ = 0.0317), while no further significant interactions were observed (1-back; β = 0.90 [-0.20, 2.01], $t$(167) = 1.61, $p$ = 0.108. In addition, the inclusion of group allocation as a covariate within the model yielded similar interaction effects (1-back: β = 0.98 [-0.11, 2.08], $t(169) = 1.77$, $p = 0.0785$; 2-back: β = 1.36 [0.24, 2.47], $t(178) = 2.40$, $p = 0.0175$; 3-back: β = 1.32 [0.22, 2.42], $t(175) = 2.36$, $p = 0.0194$).

In addition to the median-split $T_{er}$ analysis on posterior accuracy (see Results, main text), we assessed the potential influence of group allocation by including it as a covariate within the median-split $T_{er}$ analysis. Here, the significant $T_{er}$ × task complexity interaction persisted for both the observed ($F[3,149] = 2.86$, $p = 0.0391$, $\eta_p^2 = 0.05$ [0.00, 1.00]) and posterior data models ($F[3,149] = 3.05$, $p = 0.0305$, $\eta_p^2 = 0.04$ [0.00, 1.00]).

#### Supplementary Note 8

An exploratory analysis across all task trials (*n*=60) on the Probabilistic Instrumental Learning Task was undertaken. Consistent with the primary analyses, those who received pitolisant showed higher optimal choices ($F[1,55] = 7.03$, $p = 0.0104$, $\eta_p^2 = 0.11$ [0.01, 0.28]; win trials EMM = 0.13 ± 0.06, $p = 0.0285$, $d = 0.75$ [0.07, 1.43]; loss trials EMM = 0.13 ± 0.06, $p = 0.0309$, $d = 0.74$ [0.06, 1.42]), while there was no significant difference in time to choice ($F[1,55] = 3.16$, $p = 0.0811$).

To explore whether the significant group × task condition interaction on learning rate (log $\alpha$) was attributable to artefacts of early stochastic responding, we repeated the model after excluding the first 10 trials of each condition. The interaction remained significant (main effect of group: $F[1,54] = 9.36$, $p = 0.0035$, $\eta_p^2 = 0.15$ [0.03, 1.00]; group × task condition interaction: $F[1,49] = 4.41$, $p = 0.041$, $\eta_p^2 = 0.08$ [0.00, 1.00]; win trials EMM = 0.27 ± 0.29, $p = 0.35$; loss trials EMM = 1.21 ± 0.38, $p = 0.0023$, $d = 1.21$ [0.45, 1.97]). Moreover, there was no significant main effect of group or group × task condition interaction on inverse decision temperature ($\beta$; main effect of group: $F[1,47] = 0.32$, $p = 0.574$; group × task condition interaction: $F[1,26] = 1.60$, $p = 0.2169$).

An additional analysis on total money earned across allocation groups was conducted. Those who received pitolisant received more money overall (EMM(55) = 1.33 ± 0.53, p = 0.0156, $d = 0.66$ [0.12, 1.21]), with the drug group earning on average £1.33 more than the placebo group.

We investigated the potential relationship between RL computational parameters and drift rate ($v$) parameters across the *n*-back and recognition memory tasks. There was no significant main effect of *n*-back drift rate on both inverse decision temperature (win trials β = 0.50, $p = 0.0647$; loss trials β = 0.26, $p = 0.483$) and learning rate (win trials α = -0.02, $p = 0.635$; loss trials α = -0.01, $p = 0.751$). Similarly, there was no significant main effect of memory recognition drift rate (mean) on both inverse decision temperature (win trials β = 1.17, $p = 0.0887$; loss trials β = -0.39, $p = 0.646$) and learning rate (win trials α = -0.18, $p = 0.165$; loss trials α = 0.03, $p = 0.479$).

#### Supplementary Note 9

We examined potential indirect changes in cerebral blood flow across allocation groups which may have driven changes in BOLD signal across rsfMRI and task-based fMRI. In terms of global blood flow, there was no significant difference within either grey matter (ANOVA main effect of group: $F[1,50] = 0.64$, $p = 0.428$, two-tailed) or white matter across groups (ANCOVA main effect of group: $F[1,50] = 0.88$, $p = 0.353$) (Supplementary Fig. 16A – B). Similarly, regional blood flow within a whole-brain mask found no significant differences (pitolisant > placebo, peak $t[51] = 4.79$, $p = 0.1781$, TFCE-corrected; placebo > pitolisant, peak $t[51] = 4.91$, $p = 0.2171$, TFCE-corrected). Regional blood flow specifically within *a priori* task networks for memory consolidation/encoding (pitolisant > placebo, peak $t[51] = 4.054$, $p = 0.0581$, TFCE-corrected; placebo > pitolisant, peak $t[51] = 1.809$, $p = 0.921$, TFCE-corrected) and complex working memory (pitolisant > placebo, peak $t[51] = 3.472$, $p = 0.1712$, TFCE-corrected; placebo > pitolisant, peak $t[51] = 2.125$, $p = 0.906$, TFCE-corrected) were not significantly different.

#### Supplementary Note 10

The core mechanism of pitolisant is believed to be upregulation of synaptic histamine via autoreceptor blockade [21–23]. However, as with most psychoactive drugs [24], there may be ancillary effects present. Levels of nonhistaminergic monoamines remain unchanged in $H_3R$−/−mice, while there is evidence of clear increases in synaptic histamine [25–27]. Early work on pitolisant suggests it may selectively increase synaptic dopamine in the prefrontal cortex but not striatum [28]. However, in human dopaminergic neurons, $H_3R$ mRNA is low, and therefore increased synaptic dopamine may be an indirect result of synaptic histamine on co-localised monoaminergic neuron subpopulations in the prefrontal region [22,29,30]; this is consistent with the broader network effects typical of classical monoamines at the synapse [31–33]. Indeed, given the abundance of $H_3$-heteroreceptors in the region, a lack of change in striatal dopamine after pitolisant supports its preference for $H_3$-autoreceptors [34]. Further, pitolisant may induce a brief spike (~30 min) in prefrontal acetylcholine levels, however, returning to basal state (1hr post-administration) prior to the present study testing period (3hrs) [28], in addition to very small increases in serotonin (15%) selective to the striatum and not serotonin-rich prefrontal and hypothalamic regions [35,36]. Taken together, the region-selectivity of dopamine and small or brief increases in region-selective acetylcholine and serotonin, in comparison to whole-brain changes in synaptic histamine [28,37], are consistent with the preferential action of pitolisant for histamine autoreceptors. Indeed, relative to $H_3R$ agonists, pitolisant does not show intrinsic activity at one of the longest $H_3R$ isoforms, $H_3R$-453, which is presumed to function as a heteroreceptor [38,39]; moreover, compared to $H_3R$-445, inverse agonism (*e.g.,* via pitolisant) leads to larger responses in shorter and more constitutively active isoforms, $H_3R$-365 and $H_3R$-373, which are believed to function as autoreceptors [38–40]. While pitolisant shows lower binding affinity at these short isoforms, it still produces robust inverse agonism, consistent with their high constitutive activity; moreover, expression of $H_3R$ isoforms in human tissue remains incompletely resolved, so binding affinity does not directly map onto functional effect size [38]. Nevertheless, further work is required to further identify $H_3R$ isoforms in humans (>20 isoforms identified cross-species) [41], and clarify which are preferentially targeted by pitolisant at therapeutically-relevant doses.

### Supplementary Figures

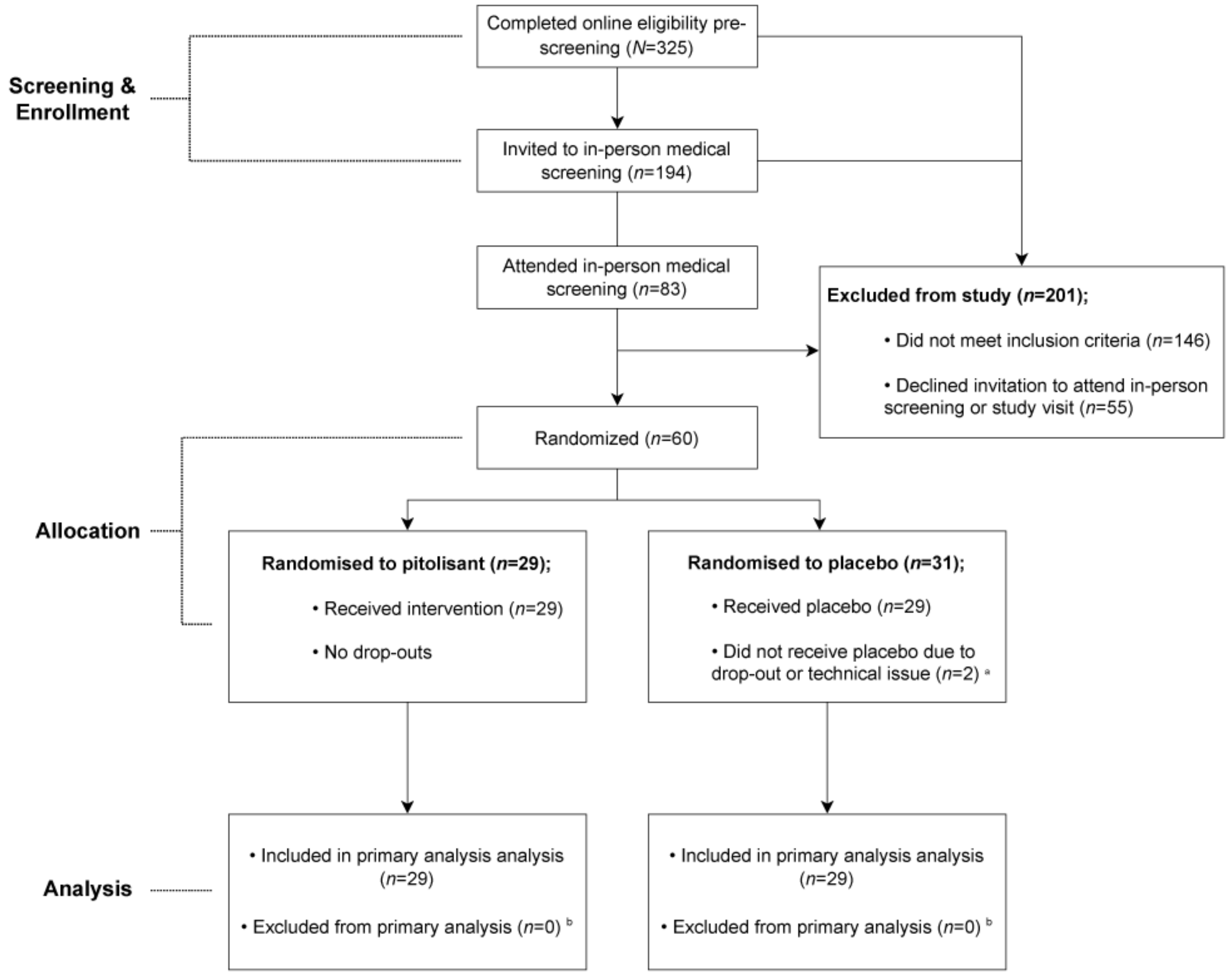

**Supplementary Fig. 1. CONSORT flowchart: recruitment and randomisation progression throughout the study.** [a] One participant randomised to placebo dropped out of the study prior to administration on the study day; another participant was withdrawn from the study due to technical issues on the study day (*i.e.,* MRI scanner coil failure).[b] Participants were included in all analyses unless there were procedure-specific data quality concerns (*e.g.,* extreme framewise displacement during fMRI tasks).

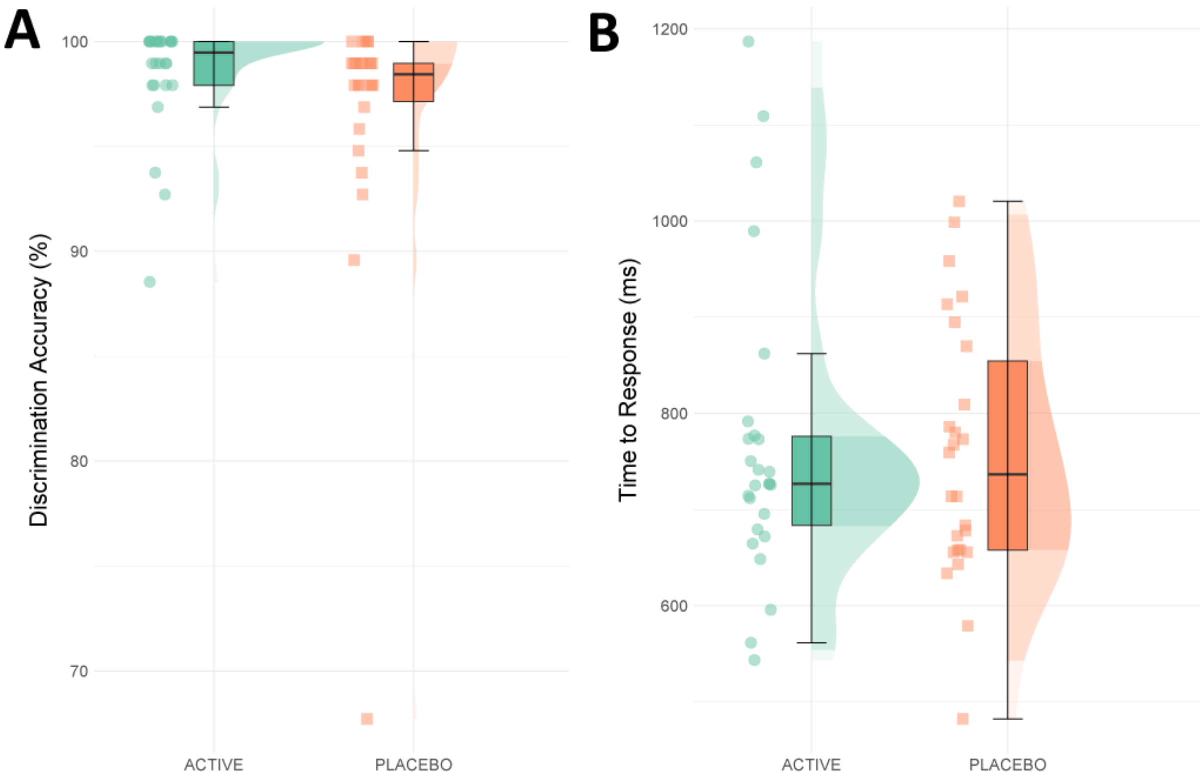

**Supplementary Fig. 2. Discrimination accuracy and response time across allocation groups during in-scanner encoding task. A** Mean percentage accuracy of discrimination between labelling stimuli ('animal' or 'landscape') across allocation groups. **B** Mean response time (ms; milliseconds) to choice during discrimination across groups. All panels contain data from N=52; lines and plot points depict mean value, with error bars and shaded areas around each line depicting standard mean error.

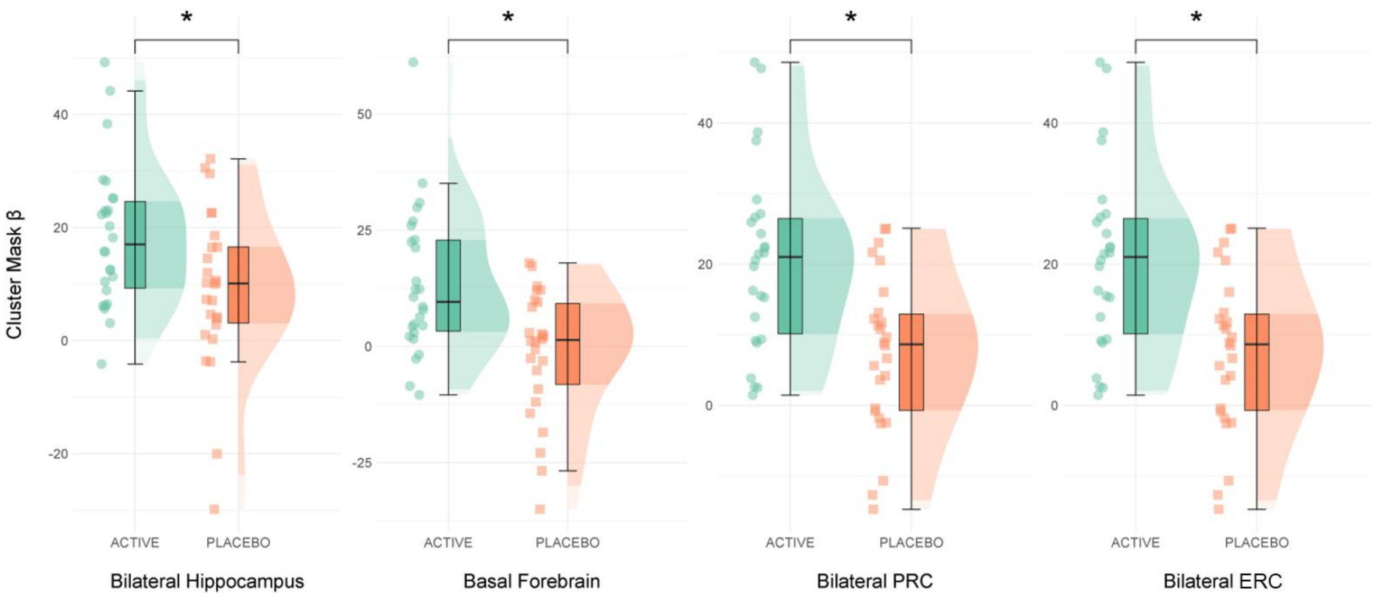

**Supplementary Fig. 3. BOLD activation (β) in a priori regions of interest during novel memory formation across allocation groups.** Parameter estimates were derived from pitolisant > placebo TFCE-corrected clusters. Boxplots represent interquartile range (IQR); central line depicts the median. Whiskers represent ±1.5 IQR, encompassing most data points; half-violin plots depict the data distribution. * $p \leq 0.05$ represents group differences by one-tailed permutation testing (TFCE corrected). Abbreviations: PRC = Perirhinal cortex; E**R**C = Entorhinal cortex.

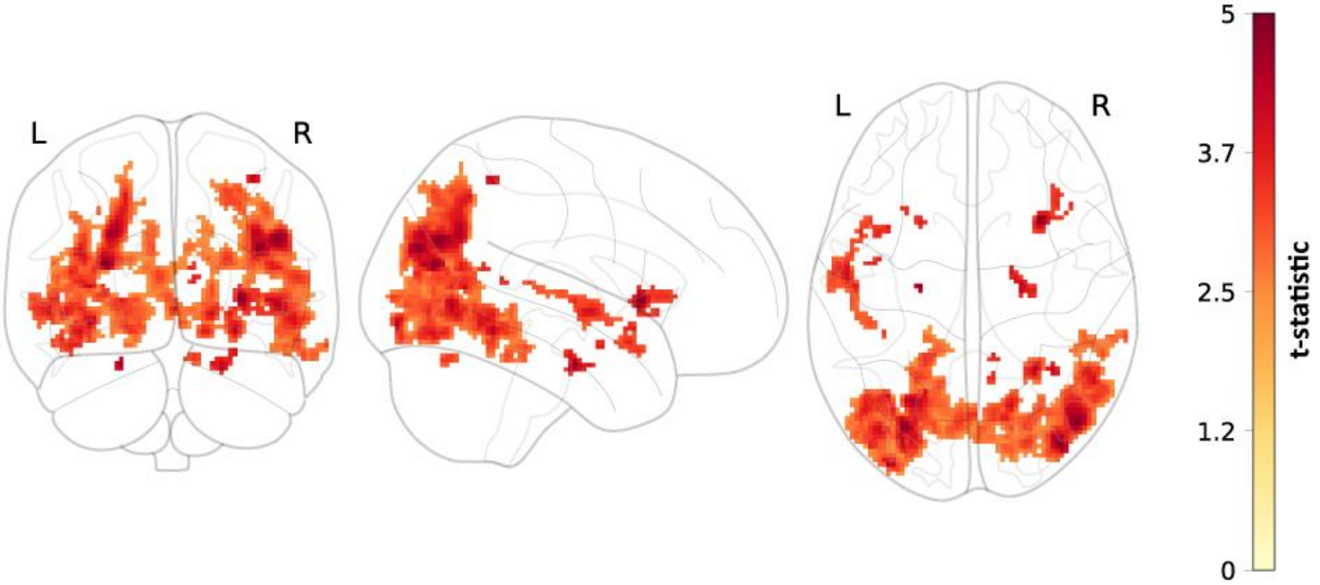

**Supplementary Fig. 4. Whole brain analysis of activity during novel memory formation across groups.** At the whole-brain level, increased activity was observed in the pitolisant group (pitolisant > placebo) during novel memory formation is represented in by the TFCE-corrected (*$p < 0.05$*) t-statistic map. The location and characteristics of each cluster is described in Supplementary Table 7.

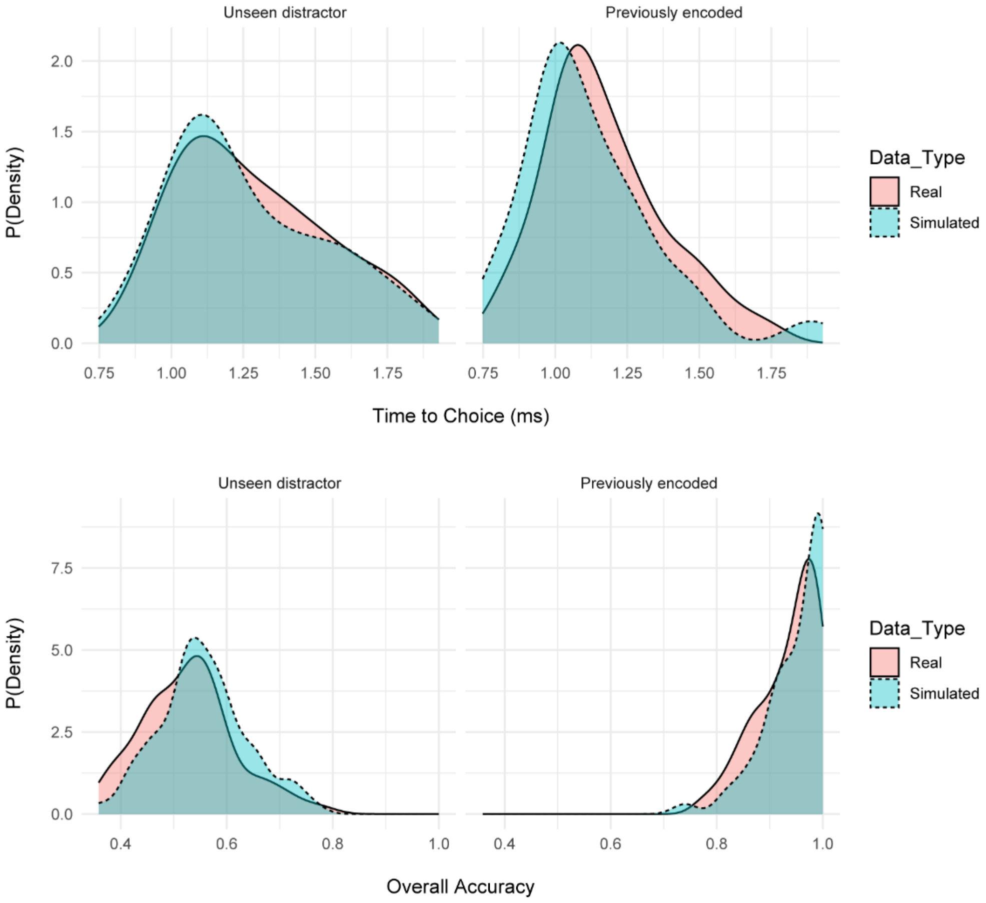

**Supplementary Fig. 5. Memory recognition DDM: Posterior predictive checks. A** Comparison of probability density curves between observed task data (red; solid line) and synthetic data generated from the model fitting process (green; dashed line), incorporating data from both study visits (baseline and follow-up). Synthetic data shown in panel A were generated from models fitted to data from N = 52 individuals (*N* = 104 total).

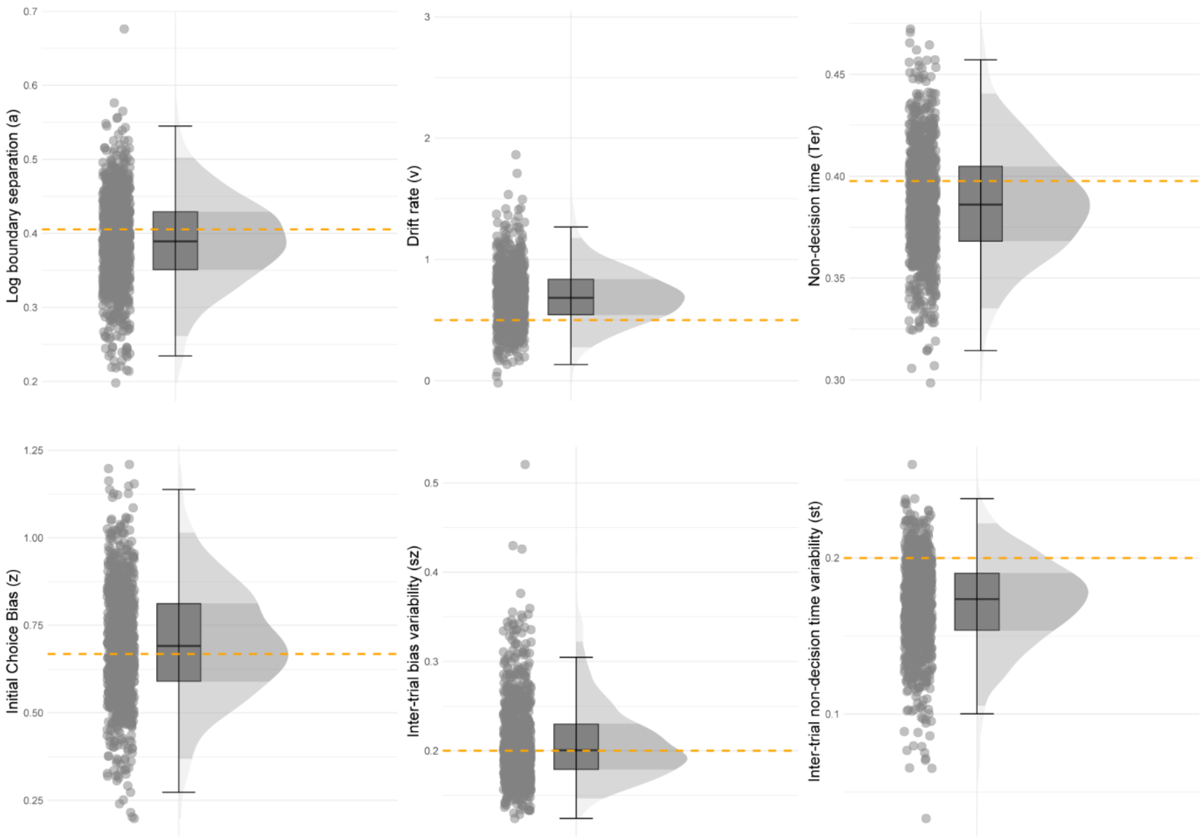

**Supplementary Fig. 6. Memory recognition DDM: Drift diffusion model parameter recovery**. Synthetic data were generated using true model parameters (yellow dashed line) and fit to the drift diffusion model. The recovered parameters from this process are plotted against their corresponding true values here. The data within figure contain 200,000 trials (or, *N*=1000 synthetic participants). Boxplots represent the interquartile range (IQR), while the central line depicts the median. The whiskers extend to approximately ± 1.5 times the IQR, encompassing the bulk of the data points; half-violin plots depict the data distribution.

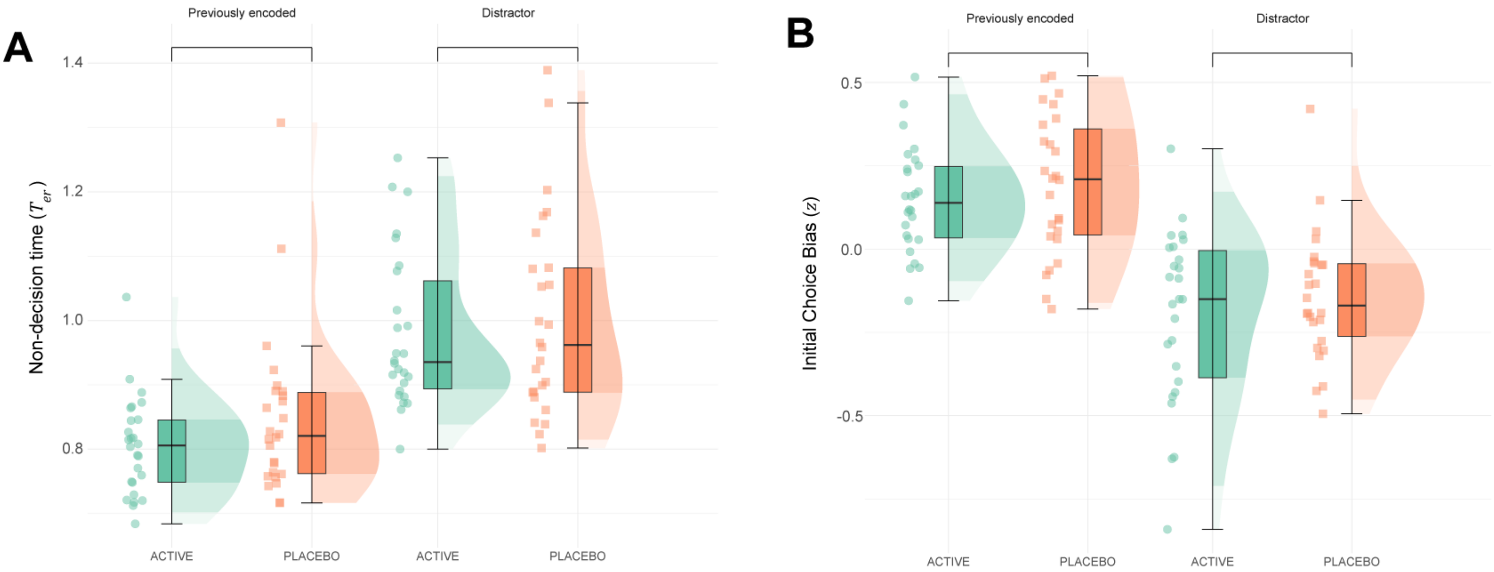

**Supplementary Fig. 7. Memory Recognition DDM: non-decision time $T_{er}$ and initial choice bias $z$ parameters across allocation groups.** There was no significant main effect of group for $\boldsymbol{T_{er}}$ (F[1,50] = 1.27, *p* = 0.266) and $z$ (F[1,50] = 1.80, *p* = 0.186). Similarly, there was no group × task condition interaction for $\boldsymbol{T_{er}}$ (F[1,50] = 0.414, *p* = 0.523) and $z$ (F[1,50] = 0.024, *p* = 0.878). All panels contain data from *N*=52; lines and plot points depict mean value, with error bars and shaded areas around each line depicting standard mean error.

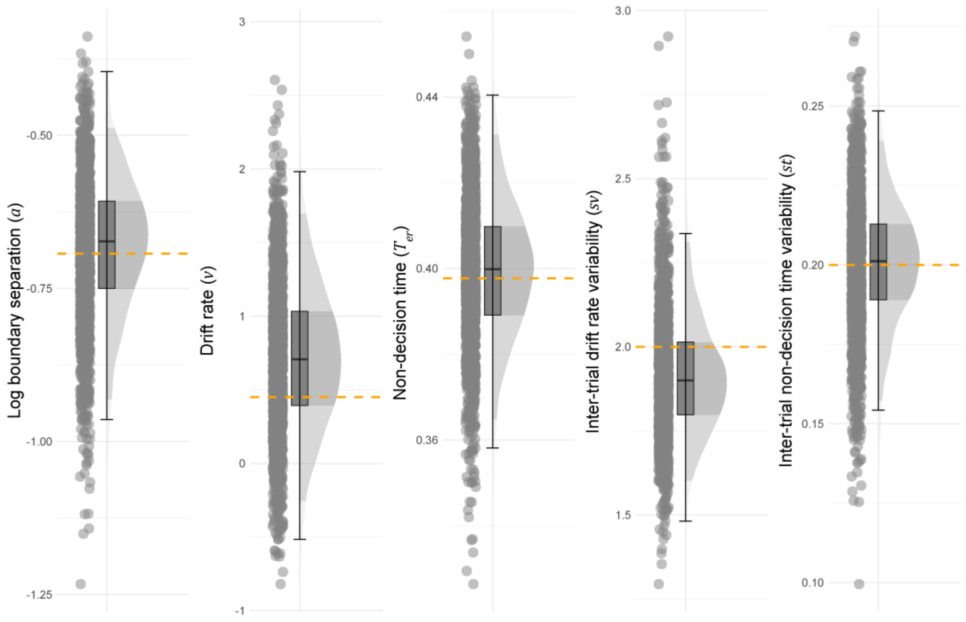

**Supplementary Fig. 8. Complex working memory task (*n*-back): Drift diffusion model parameter recovery and split non-decision time posterior predictive checks. A** Synthetic data were generated using true model parameters (yellow dashed line) and fit to the drift diffusion model. The recovered parameters from this process are plotted against their corresponding true values here. The data within figure contain 200,000 trials (or, *N*=500 synthetic participants). Boxplots represent the interquartile range (IQR), while the central line depicts the median. The whiskers extend to approximately ± 1.5 times the IQR, encompassing the bulk of the data points; half-violin plots depict the data distribution.

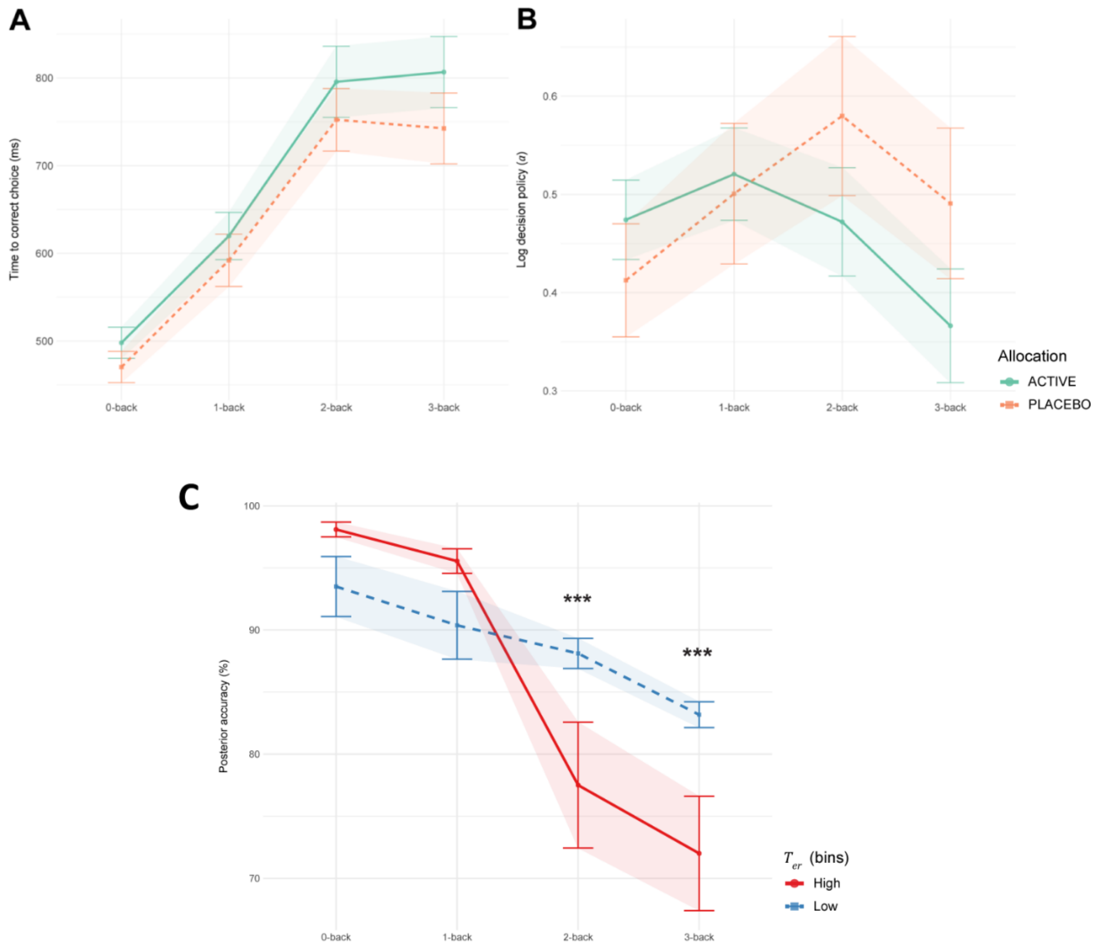

**Supplementary Fig. 9. Additional n-back analyses: time to correct choice (ms), decision policy $a$ parameter and split $T_{er}$ analysis. A** There was no significant main effect of group on time to choice (ANOVA: F[1,50] = 1.02, p = 0.317), in addition to no significant group x task condition interaction (F[3, 150] = 0.44, *p* = 0.722). **B** Similarly, there was no significant main effect of group on decision policy $a$ (F[1,50] = 0.289, *p* = 0.593), in addition to no significant group x task condition interaction (F[3,150] = 1.905, *p* = 0.131). **C** In addition to the median-split $T_{er}$ analysis on posterior accuracy (see Results, main text), we repeated the approach on observed accuracy values. This confirmed a $T_{er}$ × task complexity interaction on observed accuracy (F[3,149] = 2.857, p = 0.0391, $\eta_p^2$ = 0.13 [0.05, 0.22]). Significant pairwise differences were seen at medium-hard task difficulty, but not control and easy difficulty (0-back EMM = -0.09 ± 0.04, *p* = 0.248; 1-back EMM = 0.02 ± 0.03, *p* = 0.1552; 2-back EMM = -0.07 ± 0.03, *p* = 0.0003, *d* = -0.85 [-1.92, -0.52]; 3-back EMM = -0.10 ± 0.03, *p* = 0.0003, *d* = -1.18 [-2.12, -0.83]). All panels contain data from *N*=52; lines and plot points depict mean value, with error bars and shaded areas around each line depicting standard mean error.

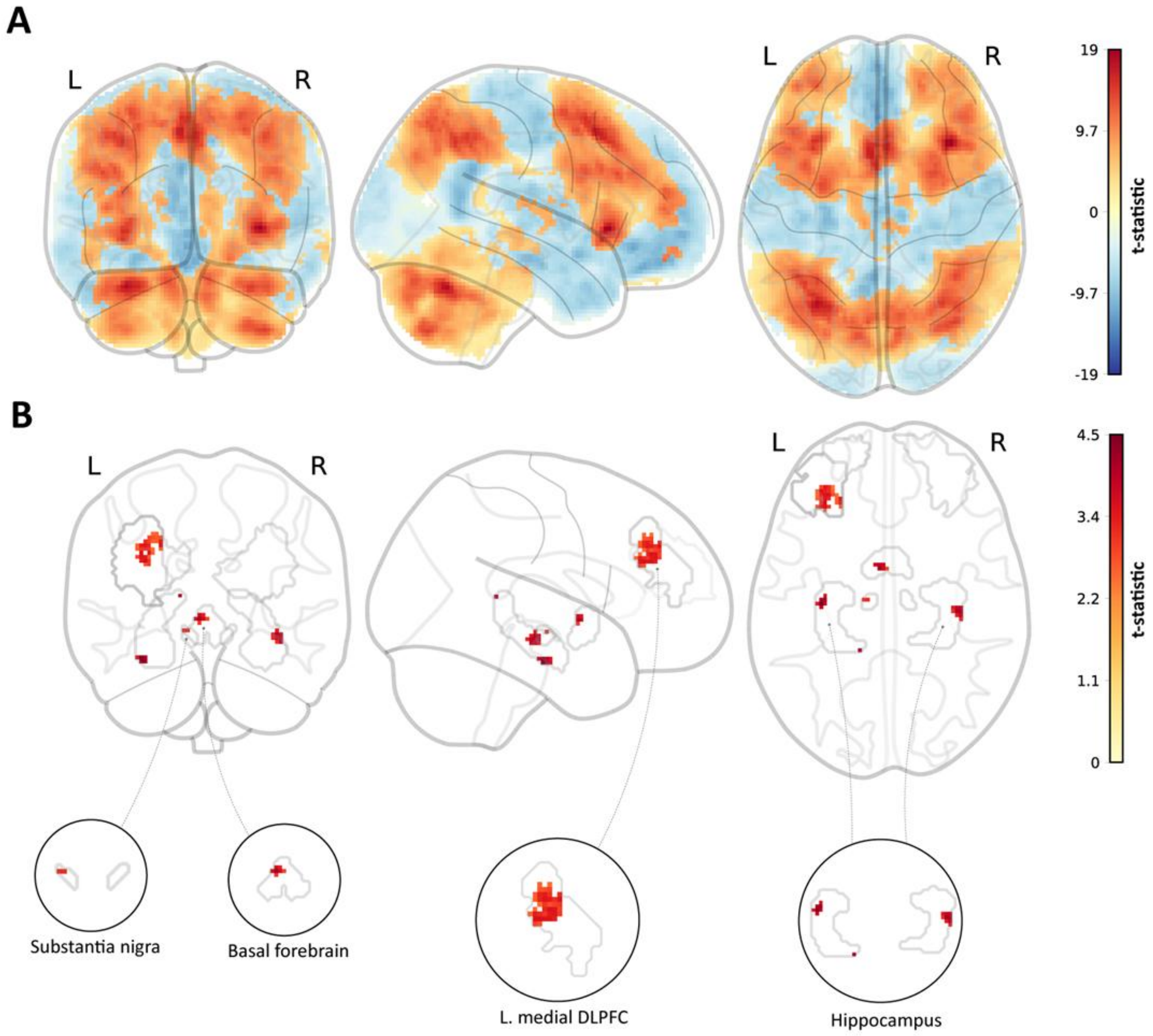

**Supplementary Fig. 10. Functional imaging analysis of the complex working memory task. A** Whole brain analysis (via permutation testing) of the effect of task is displayed for the primary contrast. TFCE-corrected t-statistic maps show increased (2 > 0-back) and decreased (0 > 2-back) BOLD activity during complex working memory processing across a network of structures including the dorsolateral prefrontal cortex, cerebellum, thalamus and brain stem (whole brain 2 > 0-back cluster: peak MNI coordinates = X,Y,Z [-6, -56, -62]; cluster size (voxels) = 79,786; t-max(51) = 3.320, p = 0.0002, TFCE-corrected; whole brain 0 > 2-back cluster 1: peak MNI coordinates = [-48, -2, -46]; cluster size = 50,848; t-max(51) = 3.810, p = 0.0002, TFCE-corrected; whole brain 0 > 2-back cluster 2: peak MNI coordinates = [-20, -104, 8]; cluster size = 1,842; t-max(51) = 6.529, p = 0.0014, TFCE-corrected; whole brain 0 > 2-back cluster 3: peak MNI coordinates = [-50, -64, 34]; cluster size = 260; t-max(51) = 6.245, p = 0.0120, TFCE-corrected; whole brain 0 > 2-back cluster 4: peak MNI coordinates = [34, -78, -34]; t-max(51) = 8.022, p = 0.0154, TFCE-corrected). **B** Increased activity during complex working memory processing (2 > 0-back) in the pitolisant group (pitolisant > placebo; SVC permutation testing) across *a priori* ROIs (significant clusters displayed in tan) including the medial DLPFC, bilateral hippocampus, basal forebrain and substantia nigra (see Supplementary Table 6 and Supplementary Fig. 7 for further information); *t*-statistics maps are TFCE-corrected. Enlarged ROIs are provided for visual purposes, with the ROI mask displayed in the gray outline. Panels A – B contain data for *N*=52 individuals. Abbreviations: BOLD = Blood Oxygenation Level Dependent; DLPFC = Dorsolateral Prefrontal Cortex; TFCE = Threshold-Free Cluster Enhancement; ROI = Region of Interest.

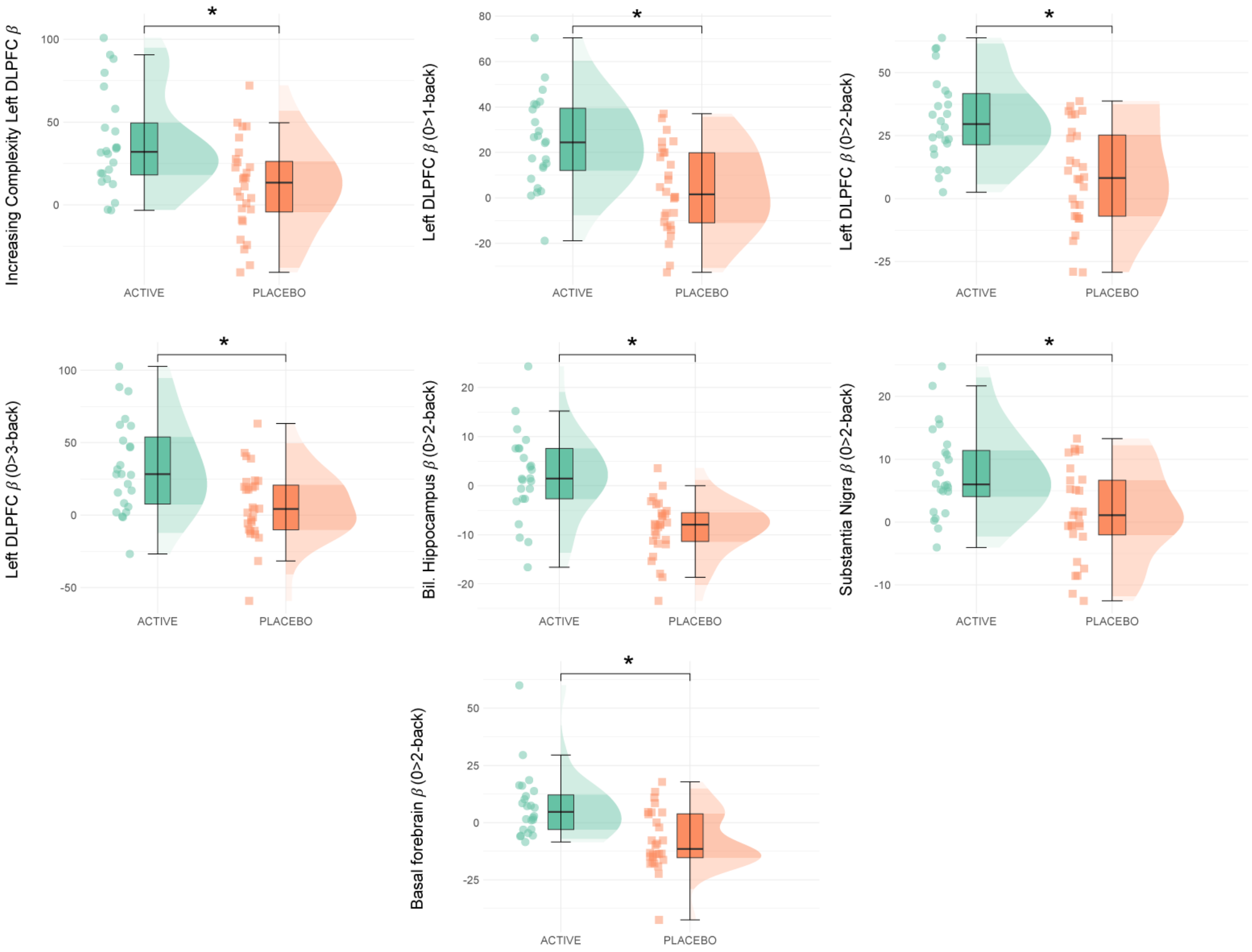

**Supplementary Fig. 11. BOLD activation (β) in a priori regions of interest during the complex working memory task across allocation groups.** Parameter estimates were derived from pitolisant > placebo TFCE-corrected clusters. Boxplots represent interquartile range (IQR); central line depicts the median. Whiskers represent ±1.5 IQR, encompassing most data points; half-violin plots depict the data distribution. * $p \leq 0.05$ represents group differences by one-tailed permutation testing (TFCE corrected). Abbreviations: DLPFC = Dorsolateral prefrontal cortex.

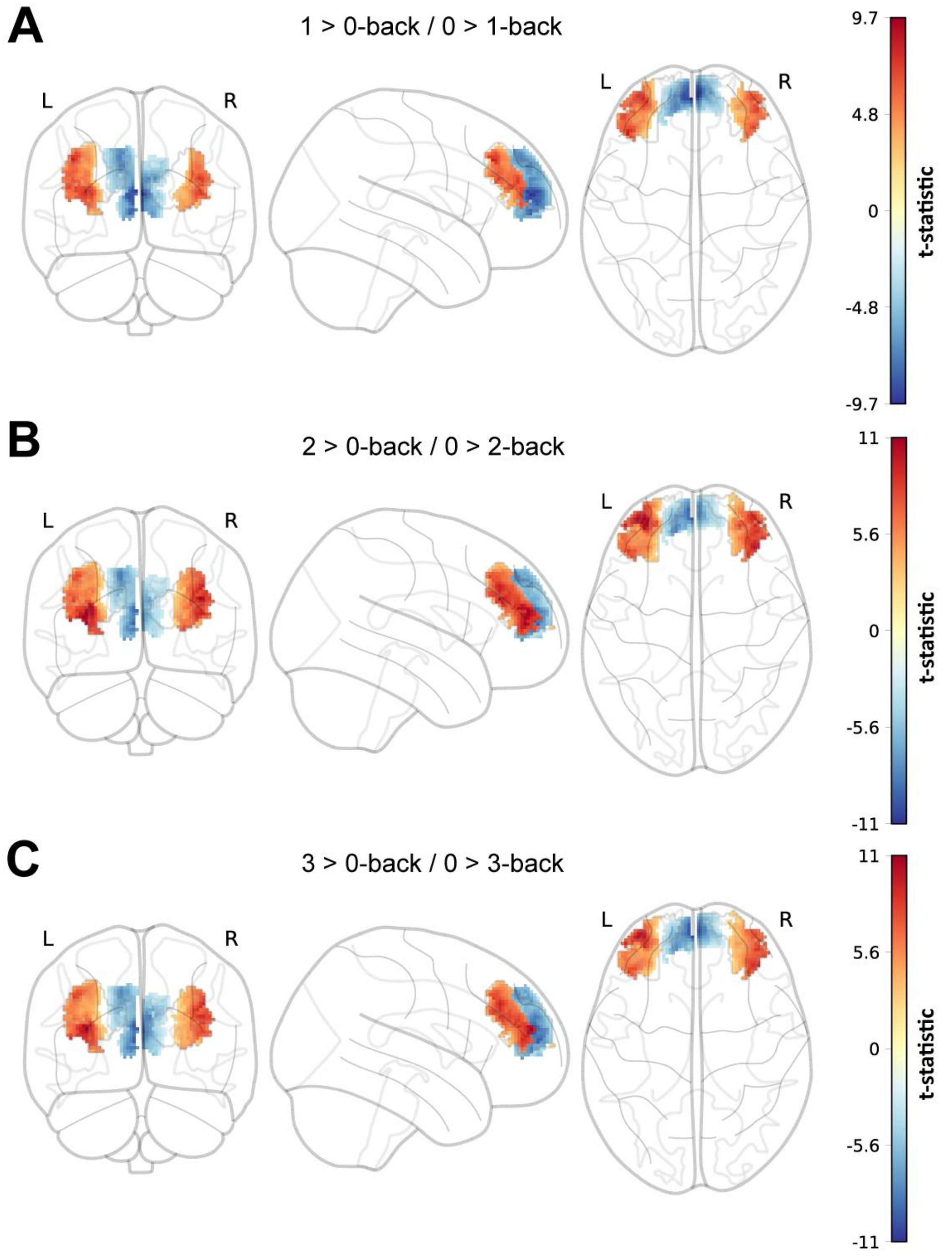

**Supplementary Fig. 12. N-Back fMRI paradigm: main effect of task across each level of task complexity.** Small volume corrected analysis of the effect of task on medial and rostral DLPFC activity across each level of task complexity (**A**: 1 > 0-back / 0 > 1-back; **B**: 2 > 0-back / 0 > 2-back; **C**: 3 > 0-back / 0 > 3-back). TFCE-corrected ($p < 0.05$) t-statistic maps show increased (1/2/3 > 0-back) and decreased (0 > 1/2/3-back) BOLD activity during complex working memory processing. Increased DLPFC peaked in the 2 > 0-back condition by number of voxels activated (3 > 0-back cluster left DLPFC cluster: peak MNI coordinates = X,Y,Z [-28, 48, 4]; cluster size = 893; $t$-max (51) = 5.280, $p < 0.001$, TFCE-corrected; 3 > 0-back cluster right DLPFC cluster: peak MNI coordinates: [28, 50, 6]; cluster size = 837; $t$-max (51) = 5.284, $p < 0.001$, TFCE-corrected; 2 > 0-back left DLPFC cluster: peak MNI coordinates = [-28, 48, 4]; cluster size = 977; $t$-max (51) = 7.252, $p < 0.001$, TFCE-corrected; 2 > 0-back right DLPFC cluster: peak MNI coordinates = [28, 50, 6]; cluster size = 871; *t*-max (51) = 8.145, $p < 0.001$, TFCE-corrected; 1 > 0-back left DLPFC cluster: peak MNI coordinates = [-32, 50, 6]; cluster size = 738; *t*-max (51) = 4.997, $p < 0.001$, TFCE-corrected; 1 > 0-back right DLPFC cluster: peak MNI coordinates = [32, 54, 8]; cluster size = 552; *t*-max (51) = 5.301, $p < 0.001$, TFCE-corrected). Further, decrease activity in the DLPFC peakjed in the 0 > 1-back condition by number of voxels activated (0 > 3-back left DLPFC cluster: peak MNI coordinates = [-6, 50, 0]; cluster size = 735; *t*-max = 6.805, $p < 0.001$, TFCE-corrected; 0 > 3-back right DLPFC cluster: peak MNI coordinates = [10, 58, 2]; cluster size = 622; *t*-max (51) = 4.539, $p < 0.001$, TFCE-corrected; 0 > 2-back left DLPFC cluster: peak MNI coordinates = [-6, 50, 0]; cluster size = 717; *t*-max (51) = 6.974, $p < 0.001$, TFCE-corrected; 0 > 2-back right DLPFC cluster: peak MNI coordinates = [2, 52, 6]; cluster size = 573; *t*-max (51) = 4.460, $p < 0.001$, TFCE-corrected; 0 > 1-back left DLPFC cluster: peak MNI coordinates = [-6, 50, 0]; cluster size = 747; *t*-max (51) = 6.064, $p < 0.001$, TFCE-corrected; 0 > 1-back right DLPFC cluster: peak MNI coordinates = [12, 52, 2]; cluster size = 672; *t*-max (51) = 4.218, $p < 0.001$, TFCE-corrected).

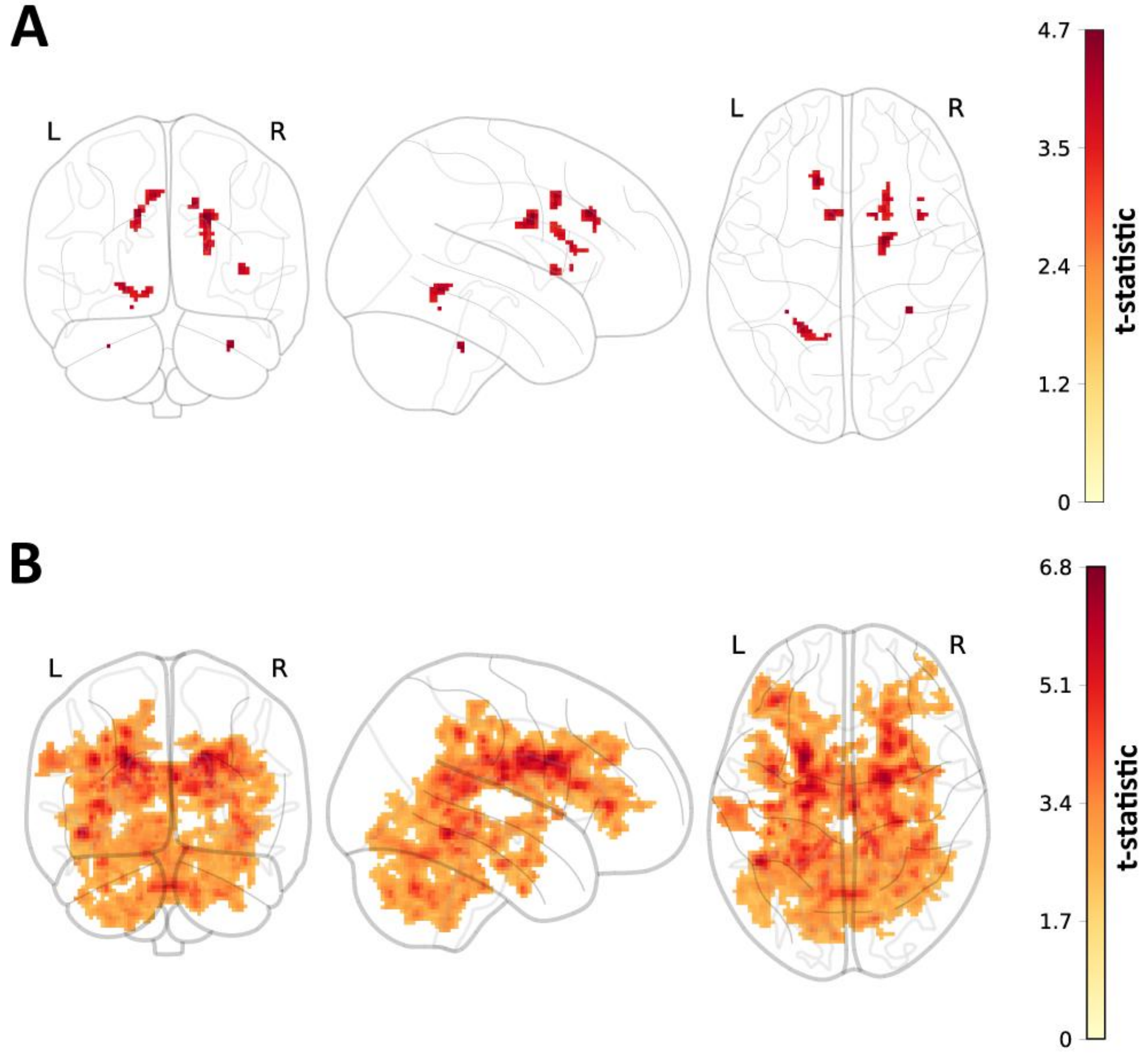

**Supplementary Fig. 13. Whole brain analysis of activity during complex working memory across groups.** At the whole-brain level, increased activity was observed in the pitolisant group (pitolisant > placebo) during during 1 > 0-back (**A**) and 2 > 0-back (**B**) is represented in by the TFCE-corrected (*p < 0.05*) t-statistic map. No significant clusters were identified in the whole-brain analysis of the 3 > 0-back contrast. The location and characteristics of each cluster is described in Supplementary Table 4.

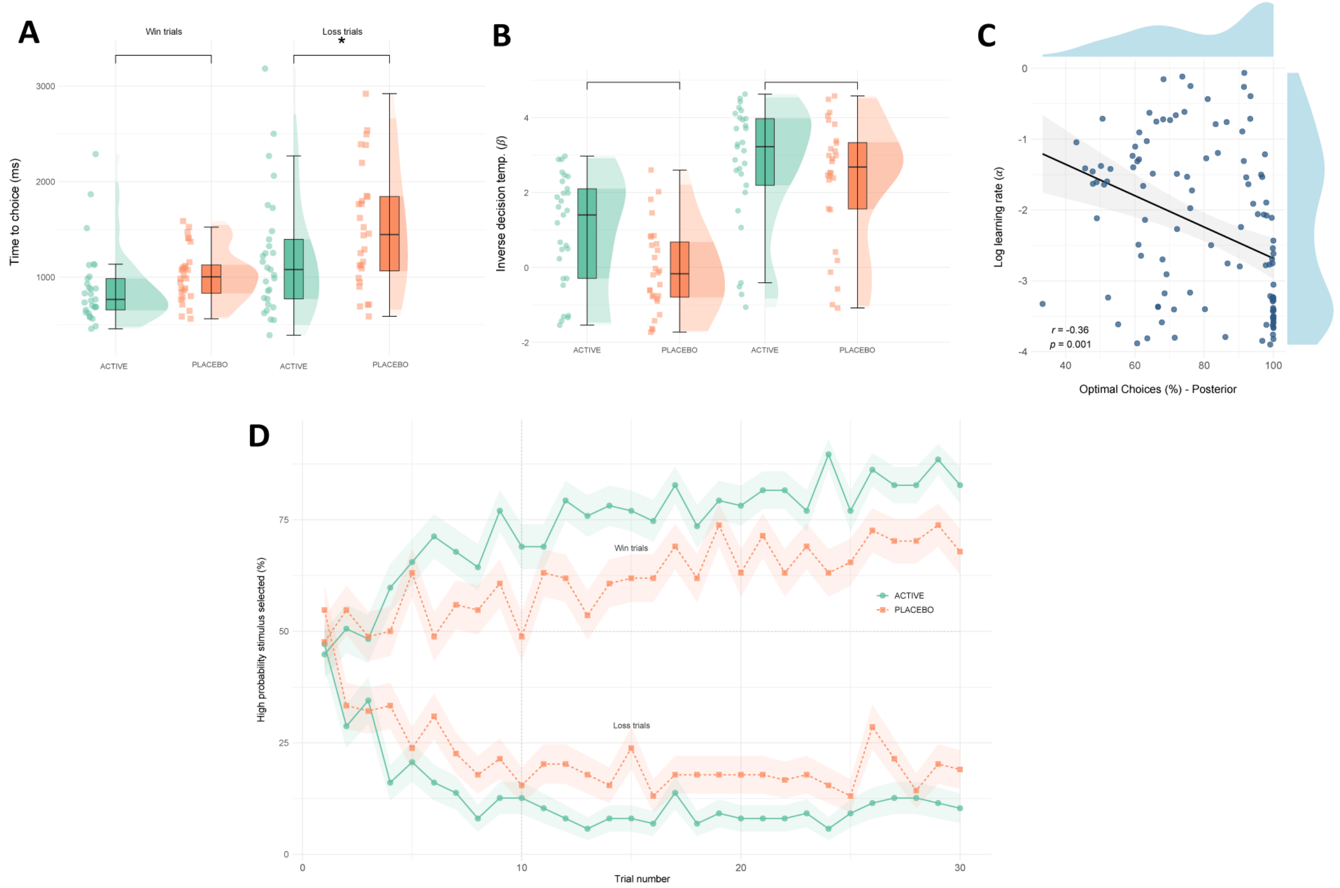

**Supplementary Fig. 14. Additional analyses - Probabilistic Instrumental Learning Task. A** There was no significant main effect of group on time to choice (ms) during the task (F[1,55] = 3.263, *p* = 0.0763), in addition to no significant group × task condition interaction (F[1,55] = 2.02, *p* = 0.16). **B** Similarly, there was no significant effect of group on inverse decision temperature during the task (F[1,42] = 2.40, *p* = 0.1287), in addition to no significant group × task condition interaction (F[1,52] = 0.35, *p* = 0.5580). Boxplots represent the interquartile range (IQR), while the central line depicts the median. The whiskers extend to approximately ± 1.5 times the IQR, encompassing the bulk of the data points; half-violin plots depict the data distribution. C Model-fitted posterior for optimal choices were associated with learning rate (β = -5.55, *p* = 0.00142, $\eta_p^2$ = 0.16 [0.07, 1.00]; *r* = -0.36); this relationship was observable within allocation groups, based on aggregated posterior estimates (β = -5.09 [-7.00, -3.19]). The fitted line through data points was generated via linear modelling (shaded area: standard mean error). **D** Synthetic temporal learning was generated using posterior choice probabilities and simulating choices from a Bernoulli distribution. Synthetic learning over time across groups (averaged across blocks) are displayed here – see Figure 5B (main text) for comparison with observed data. The Y axis (‘high probability stimulus selected’) refers to the selection (mean %) of high probability win or loss symbols. Plot points depict mean value; error bars and shaded area depict SEM. Panels **A**–**D** includes data from *N*=57 individuals.

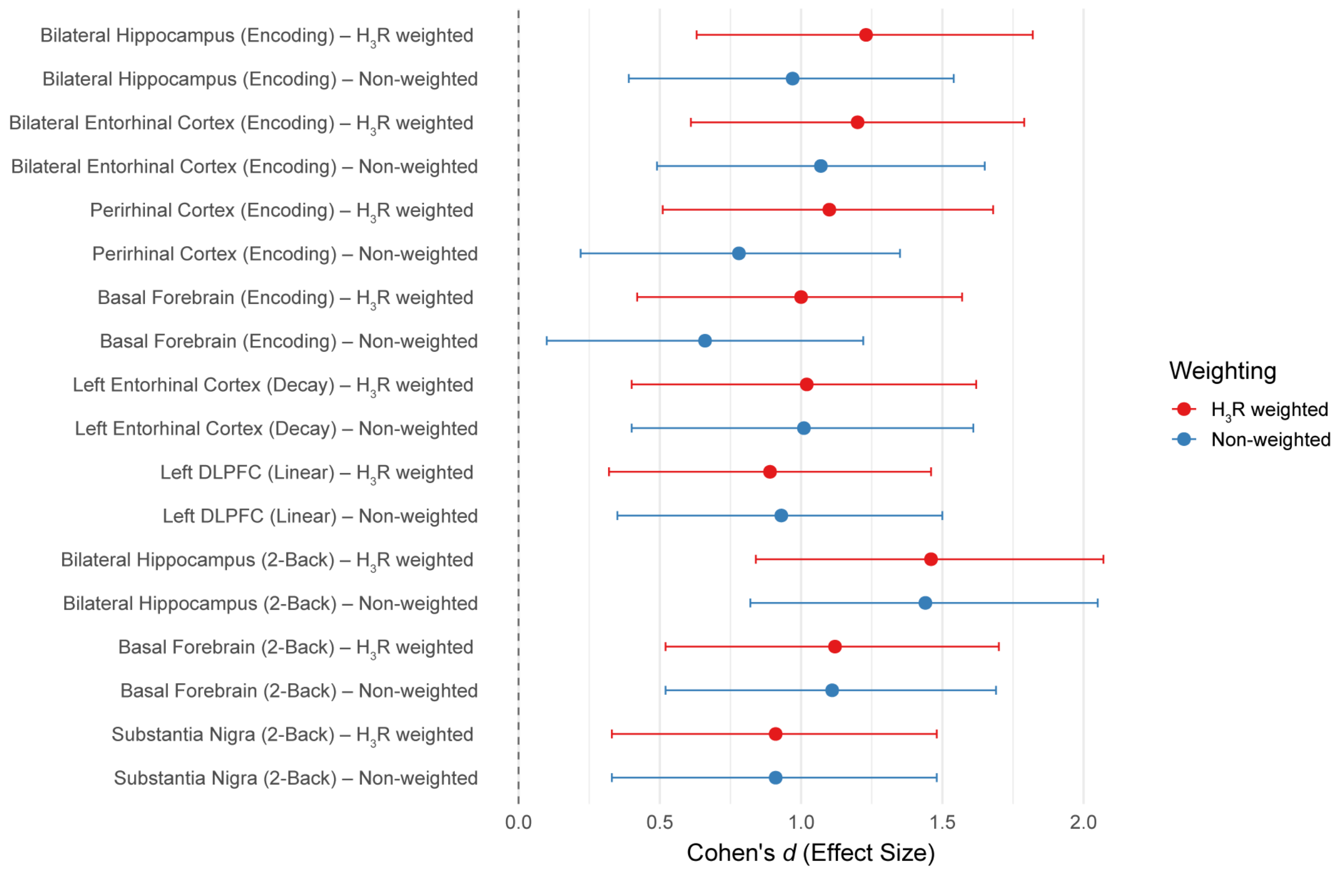

**Supplementary Fig. 15. Summary of fMRI task-based group effect cluster effect sizes: $H_3R$-weighting vs non-weighting.** Throughout the fMRI task-based analysis (memory encoding and n-back fMRI), multiple group-based effects (drug > placebo) were identified via TFCE-corrected permutation testing. BOLD signal within the cluster (alpha thresholded; $p < 0.05$) was extracted within using both binary and $H_3R$-weighted cluster masks. Central line dots represent mean Cohen's d, and whiskers represent the 95% confidence interval.

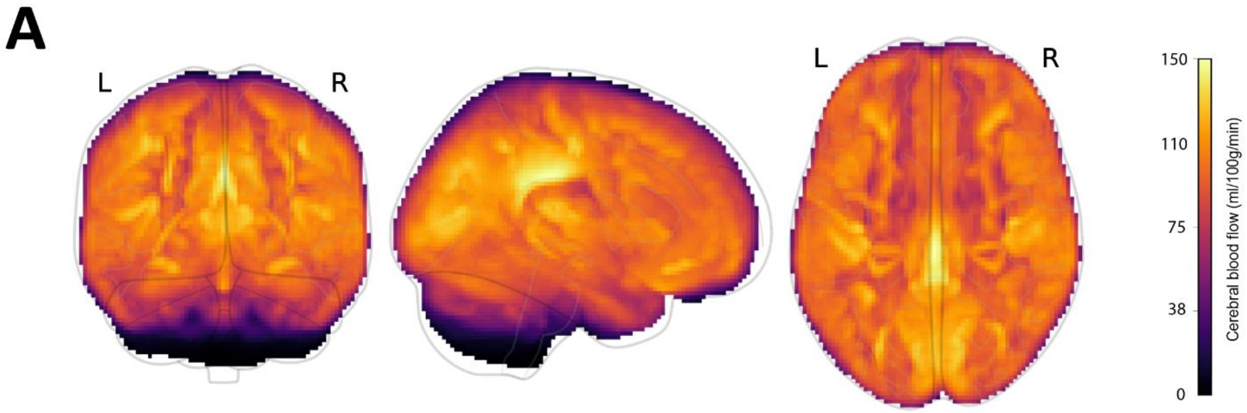

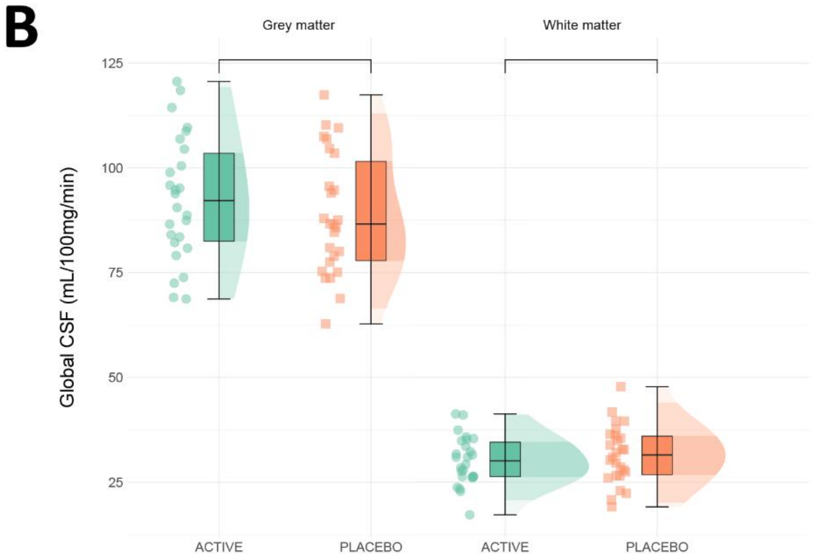

**Supplementary Fig. 16. Arterial Spin Labelling – Cerebral Blood Flow Analysis. A** Mean Cerebral blood flow across all participants. **B** Global cerebral blood flow values (grey and white matter) **across allocation groups.** All panels contain data from *N*=52; boxplots represent the interquartile range (IQR), while the central line depicts the median. The whiskers extend to approximately ± 1.5 times the IQR, encompassing the bulk of the data points; half-violin plots depict the data distribution.

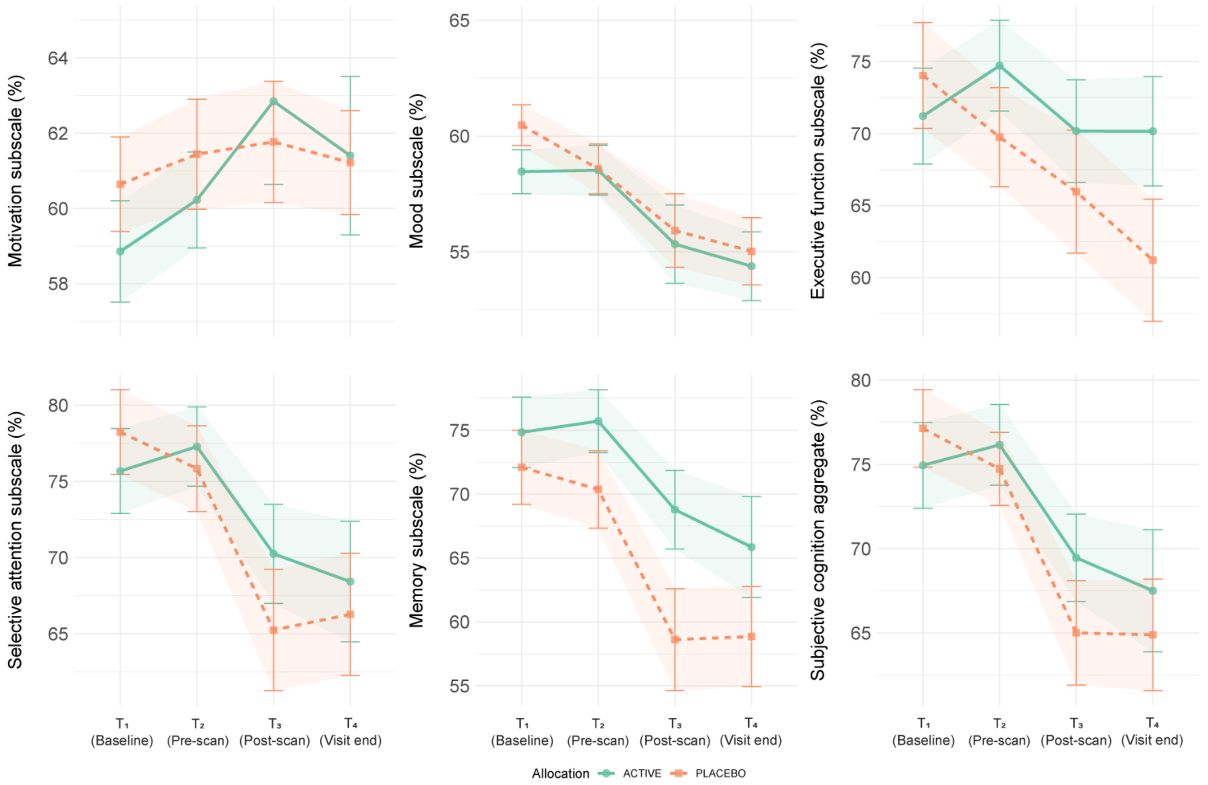

**Supplementary Fig. 17. Subjective cognition throughout the acute dosing period across allocation groups.** Throughout the acute dosing period, individuals were asked to self-rate subjective cognition at regular intervals ($T_1$: Baseline, before intervention; $T_2$ pre-scan, 2.5hrs post-intervention; $T_3$ post-scan, 4hrs post-intervention; $T_4$ at visit end, 5hrs post-intervention). Scores for each allocation group are plotted across all timepoints, where higher scores indicate greater self-assessment of cognitive ability, and were divided into five subscales: motivation, mood, executive functioning, selective attention, and memory. Lines and plot points depict mean value, with error bars and shaded areas around each line depicting standard mean error. This figure contains data for *N*=57 individuals (216 observations total). *N*=1 individuals were excluded from analyses due to data missingness (baseline).

#### Supplementary Tables

**Supplementary Table 1.** Study inclusion and exclusion eligibility criteria

| |
|---|
| Inclusion criteria:<br>• Participant is willing and able to give informed consent for participation in the research<br>• Not currently taking any medications which may interfere with pitolisant, including psychoactive medications<br>• Not currently using antihistaminergic medication<br>• Aged 18-45 years<br>• Male or female<br>• Sufficiently fluent English to understand and complete cognitive tasks and questionnaires<br>• Body Mass Index above or below 18-30<br>• Right handed |
| Exclusion criteria:<br>• Current pregnancy (as determined by urine pregnancy test taken during screening visit), planning to become pregnant or breast feeding<br>• Any past or current history of severe and/or serious psychiatric disorder, including but not limited to schizophrenia, psychosis, bipolar affective disorder, major depressive disorder, obsessive compulsive disorder<br>• Clinically significant abnormal values for urine drug screen, blood pressure measurement (in accordance with AP20 'non-invasive blood pressure'). A participant with a clinical abnormality or parameters outside the reference range for the population being studied may be included only if the Investigator considers that the finding is unlikely to introduce additional risk factors and will not interfere with the study procedures<br>• History of, or current medical conditions which, in the opinion of the investigator, may interfere with the safety of the participant or the scientific integrity of the study, including epilepsy/seizures, brain injury, hepatic or renal disease, acid-related gastro-intestinal problems, Central Nervous System (CNS) tumours, neurological conditions<br>• Current or past history of drug or alcohol dependency<br>• Severe lactose intolerance<br>• Use of recreational drugs (e.g. cannabis, cocaine, amphetamines) within past 3 months<br>• Participation in a study which uses the same computer tasks as those in the present study (determined by asking participants about previous studies participated in during screening) within past 3 months<br>• Participation in a study that involves the use of a medication within the last three months<br>• Smoking > 5 cigarettes per day<br>• Consumption of a high amount of caffeine per day (> 400ml caffeine) (*e.g.,* 5 or more cups of coffee)<br>• Participant is unlikely to comply with the clinical study protocol or is unsuitable for any other reason, in the opinion of the Investigator<br>• Any contraindication to MRI scanning (e.g. metal objects in your body, pacemakers, significant claustrophobia) |

**Table 2. Demographic characteristics across allocation groups**

| | Pitolisant (*n*=29) | Placebo (*n*=29) | Inferential analysis [a] |
|---|---|---|---|
| Age (Years), M (S.D.) | 27.14 (5.57) | 29.21 (6.72) | 0.21 |
| Gender, *N* male:female | 10:19 | 11:18 | 1.00 |
| Body mass index, M (S.D.) | 23.90 (3.49) | 23.59 (3.23) | 0.73 |
| Contraceptive use, yes:no | 6:13 | 5:13 | 1.00 |
| Native language, *N* | | | 0.67 |
| English | 21 | 23 | 0.50 |
| Chinese | 2 | 1 | |
| Spanish | 1 | 3 | |
| Other | 5 | 2 | |
| Digit span aggregate | 36.83 (5.67) | 36.10 (6.10) | 0.64 |
| Time in education (Years), M (S.D.) | 17.17 (2.82) | 17.97 (2.64) | 0.27 |
| Highest educational attainment, *N* | | | 0.71 |
| High School / Sixth form | 5 | 6 | |
| Undergraduate degree | 12 | 8 | |
| Postgraduate (Masters) | 8 | 11 | |
| Postgraduate (Doctorate) | 4 | 4 | |

[a] Values represent significance values inferential analyses. These values pertain to Welch's two Sample t-test where differences between group means were analysed, and Pearson's Chi-squared where differences between frequency/ratio distributions were analysed.

**Supplementary Table 3. Memory encoding fMRI task: TFCE-corrected cluster summary for main effect of task.**

| Task contrast | Cluster | Location(s)* | Hemisphere / division (peak) | Peak voxel coordinates (MNI) | | | Cluster size (voxels) | *t*-value (peak) | *p*-value (peak) |
|---|---|---|---|---|---|---|---|---|---|
| | | | | X | Y | Z | | | |
| Novel > Familiar | | | | | | | | | |
| | 1 | Cerebellum; Bilateral hippocampus; bilateral putamen; bilateral occipital cortex; bilateral amygdala; bilateral thalamus; bilateral pallidum; bilateral precentral gyrus; brain stem; bilateral frontal poles; bilateral Inferior frontal gyrus | Left | -32 | -70 | -54 | 62965 | 3.773 | 0.0002 |
| | 2 | Left premotor cortex | Left | -4 | 8 | 62 | 455 | 6.365 | 0.0100 |
| | 3 | Bilateral anterior cingulate cortex | Right | 6 | 4 | 28 | 123 | 5.985 | 0.0206 |
| Familiar > Novel | | | | | | | | | |
| | 1 | Bilateral precuneous cortex | Right | 12 | -64 | 26 | 6380 | 6.522 | 0.0002 |
| | 2 | Lateral Occipital Cortex; angular gyrus | Right | 54 | -62 | 40 | 1198 | 6.125 | 0.0002 |
| | 3 | Frontal pole | Right | 14 | 54 | -2 | 128 | 4.742 | 0.0002 |
| | 4 | Frontal pole | Left | -16 | 94 | 37 | 60 | 5.950 | 0.0002 |
| | 5 | Middle frontal gyrus | Right | 36 | 26 | 44 | 12 | 5.299 | 0.0020 |

Note: Only clusters identified through permutation testing which remained significant following TFCE correction ($p < 0.05$) were included. *Cortical or subcortical regions falling within the significant cluster.

**Supplementary Table 4.** Memory encoding task: Between-groups whole-brain fMRI analysis

| Task contrast | Cluster | Location(s) | Hemisphere / division (peak) | Peak voxel coordinates (MNI) | | | Cluster size (voxels) | *t*-value (peak) | *p*-value (peak) |
|---|---|---|---|---|---|---|---|---|---|
| | | | | X | Y | Z | | | |
| Novel > Familiar | | | | | | | | | |
| | 1 | Lateral occipital cortex (superior); Visual cortex (V7) | Right | 46 | -74 | 28 | 4563 | 4.957 | 0.0160 |
| | 2 | Superior, temporal and inferior temporal gyri | Left | -54 | -8 | -2 | 180 | 4.002 | 0.0400 |
| | 3 | Lingual and fusiform gyri | Right | 18 | -66 | -4 | 131 | 3.981 | 0.0408 |
| | 4 | Fusiform gyrus | Right | 28 | -50 | -10 | 101 | 4.291 | 0.0366 |
| | 5 | Insular cortex | Right | 30 | 16 | 2 | 92 | 4.825 | 0.0346 |
| | 6 | Hippocampus; parahippocampal gyrus (anterior); subiculum; entorhinal cortex | Right | 22 | -16 | -28 | 40 | 4.256 | 0.0432 |
| | 7 | Temporal pole | Left | -48 | 10 | -16 | 33 | 3.623 | 0.0470 |
| | 8 | Cerebellum (VI) | Right | 10 | -74 | -24 | 25 | 2.518 | 0.0524 |
| | 9 | Insular cortex | Left | -36 | 12 | -10 | 16 | 3.721 | 0.0476 |
| | 10 | Lateral occipital cortex | Left | -40 | -72 | 36 | 13 | 2.808 | 0.0494 |
| | 11 | Superior parietal lobule | Right | 36 | -50 | 54 | 11 | 4.089 | 0.0470 |
| | 12 | Planum polare | Left | -42 | -4 | -18 | 9 | 3.643 | 0.0488 |
| | 13 | Putamen | Left | -22 | 14 | 2 | 9 | 3.745 | 0.0488 |
| | 14 | Hippocampus; subiculum | Left | -22 | -14 | -26 | 7 | 4.651 | 0.0470 |
| | 15 | Posterior cingulate gyrus | Right | 8 | -54 | 16 | 6 | 3.594 | 0.0488 |
| | 16 | Insular cortex | Left | -28 | 18 | 4 | 5 | 3.421 | 0.0500 |
| | 17 | Posterior cingulate gyrus | Right | 10 | -46 | 12 | 5 | 3.667 | 0.0486 |
| | 18 | Lateral occipital cortex (superior) | Left | -32 | -78 | 40 | 3 | 2.929 | 0.0496 |
| | 19 | Planum temporale | Left | -38 | -34 | 10 | 2 | 3.328 | 0.0496 |
| Familiar > Novel | | | | | | | | | |
| | *NA* | -- | -- | -- | -- | -- | -- | -- | -- |

Note: Only clusters identified through between-groups permutation tests which remained significant following TFCE correction ($p < 0.05$) were included.

**Supplementary Table 5**. Post-encoding memory recognition task: Drift Diffusion Model comparison

| Model specification | Deviance information criterion (DIC) |
|---|---|
| $a + v(single) + T_{er}$ | 10457.33 |
| $a + v(stim) + T_{er}$ | 9131.99 |
| $a(stim) + v(stim) + T_{er}(single)$ | 9114.83 |
| $a(stim) + v(stim) + T_{er}(stim)$ | 8722.27 |
| $a(stim) + v(stim) + T_{er}(stim) + z(single)$ | 8722.14 |
| $a(stim) + v(stim) + T_{er}(stim) + z(stim)$ | 8721.08 |
| $a(stim) + v(stim) + T_{er}(stim) + z(stim) + st$ | 8632.29 |
| $a(stim) + v(stim) + T_{er}(stim) + z(stim) + sz$ | 8690.91 |
| $a(stim) + v(stim) + T_{er}(stim) + z(stim) + sv$ | 8689.85 |
| $a(stim) + v(stim) + T_{er}(stim) + z(stim) + sv + st$ | 8633.83 |
| $a(stim) + v(stim) + T_{er}(stim) + z(stim) + sv + sz$ | 8691.04 |
| $a(stim) + v(stim) + T_{er}(stim) + z(stim) + st + sz$ | 8631.22 |
| $a(stim) + v(stim) + T_{er}(stim) + z(stim) + sv + st + sz$ | 8635.84 |

**Note**: ‘single’ and ‘stim’ refer to parameters which are fixed or varying according to stimulus type (‘previously encoded’ or ‘distractor’ recognition images), respectively.

**Supplementary Table 6. N-Back fMRI between-groups analysis: small volume TFCE-corrected summary of a priori regions of interest across individual task contrasts (Drug > Placebo).**

| Task contrast | Mask [a] | Cluster | Hemisphere / division [b] | Peak voxel coordinates (MNI) | | | Cluster size (voxels) | *t*-value (peak) | *p*-value (peak) |
|---|---|---|---|---|---|---|---|---|---|
| | | | | X | Y | Z | | | |
| 1 > 0-Back | | | | | | | | | |
| | DLPFC (medial) | 1 | Left | -34 | 38 | 30 | 12 | 3.681 | 0.0450 |
| 2 > 0-Back (primary) | | | | | | | | | |
| | DLPFC (medial) | 1 | Left | -32 | 34 | 26 | 77 | 3.585 | 0.0124 |
| | Hippocampus | 1 | Right | 38 | -22 | -14 | 20 | 4.059 | 0.0184 |
| | | 2 | Left | -34 | -18 | -26 | 12 | 4.465 | 0.0170 |
| | | 3 | Left | -14 | -42 | 8 | 1 | 4.065 | 0.0486 |
| | Substantia nigra | 1 | Left | -10 | -16 | -10 | 2 | 3.169 | 0.0254 |
| | Basal forebrain | 1 | Left | -4 | 0 | -4 | 11 | 4.106 | 0.0118 |
| 3 > 0-Back | | | | | | | | | |
| | DLPFC (medial) | 1 | Left | -34 | 40 | 30 | 7 | 3.150 | 0.0458 |

[a] *a priori* small-volume corrected masks were bilateral and/or contained all divisions. [b] Only clusters identified through between-groups permutation tests which remained significant following TFCE correction ($p < 0.05$) were included. Note: Table contains data for N=52 individuals.

**Supplementary Table 7.** Verbal working memory n-back task: between-groups whole-brain fMRI analysis

| Task contrast | Cluster | Location(s) * | Hemisphere / division (peak) | Peak voxel coordinates (MNI) | | | Cluster size (voxels) | *t*-value (peak) | *p*-value (peak) |
|---|---|---|---|---|---|---|---|---|---|
| | | | | X | Y | Z | | | |
| 1 > 0-Back | | | | | | | | | |
| | 1 | Caudate | Right | 20 | -2 | 32 | 53 | 4.698 | 0.0382 |
| | 2 | Caudate | Right | 22 | 18 | 16 | 45 | 4.157 | 0.0448 |
| | 3 | Lingual gyrus | Left | -22 | -48 | -6 | 43 | 4.421 | 0.0438 |
| | 4 | Cingulate sulcus | Left | -14 | 28 | 34 | 37 | 4.730 | 0.0410 |
| | 5 | Paracingulate gyrus | Left | -6 | 10 | 42 | 27 | 4.373 | 0.0440 |
| | 6 | Insular cortex | Right | 40 | 10 | 4 | 15 | 4.061 | 0.0468 |
| | 7 | Paracingulate gyrus | Right | 16 | 10 | 38 | 9 | 4.365 | 0.0458 |
| | 8 | Cerebellum (VI) | Right | 32 | -38 | -36 | 8 | 4.597 | 0.0472 |
| | 9 | Frontal operculum | Right | 38 | 18 | 6 | 4 | 3.909 | 0.0492 |
| | 10 | Cerebellum (VI) | Left | -30 | -40 | -36 | 1 | 4.289 | 0.0498 |
| | 11 | Cerebellum (V) | Left | -18 | -50 | -16 | 1 | 4.102 | 0.0496 |
| 2 > 0-Back (primary) | | | | | | | | | |
| | 1 | Left medial DLPFC; bilateral hippocampus; right inferior frontal gyrus; bilateral frontal pole; bilateral caudate; bilateral thalamus; bilateral cerebellum; left substantia nigra; left pallidum; basal forebrain; left lingual gyrus; left stria terminalis; left precentral gyrus | Right | -12 | -10 | 22 | 18421 | 5.034 | 0.0006 |
| | 2 | Postcentral and anterior supramarginal gyri | Left | -60 | -20 | 26 | 298 | 4.049 | 0.0282 |
| | 3 | Insular cortex | Right | 36 | 10 | 0 | 38 | 3.782 | 0.0422 |
| | 4 | Inferior temporal gyrus | Left | -48 | -30 | -26 | 3 | 3.516 | 0.0494 |
| 3 > 0-Back | | | | | | | | | |
| | *NA* | -- | -- | -- | -- | -- | -- | -- | -- |

Note: Only clusters identified through between-groups permutation tests which remained significant following TFCE correction ($p < 0.05$) were included. *Cortical or subcortical regions falling within the significant cluster.

**Supplementary Table 8**. *n*-Back Task: Drift Diffusion Model comparison

| Model specification | Deviance information criterion (DIC) |
|---|---|
| $a + v(single) + T_{er}$ | 5766.366 |
| $a + v(split) + T_{er}$ | 1622.017 |
| $a + v(split) + T_{er} + z$ | 1513.519 |
| $a + v(single) + T_{er} + z(split)$ | 5588.073 |
| $a + v(split) + T_{er} + trial\ var.$ | 1561.630 |
| $a + v(split) + T_{er} + trial\ var. cond$[a] | 1402.282 |

[a] trial variation parameters depending on task condition (*n*-back level).

**Supplementary Table 9.** Comparison between $H_3R$-weighted and unweighted associations between group-level effects

| Relationship | Weighting | $\beta$ | *p*-corr [1] | $\eta_p^2$ [95% CI] | *r* |
|---|---|---|---|---|---|
| Hippocampus Encoding $\beta$ ~ Mam ↔ Hippocampus Connectivity | $H_3R$-weighted [2] | 1.20 | 0.012 | 0.16 [0.01, 0.35] | 0.40 |
| | Non-weighted | 0.68 | 0.10 | 0.08 [0.00, 0.26] | 0.29 |
| Trace persistence $\beta$ ~ Mam ↔ Hippocampus Connectivity | $H_3R$-weighted | 1.98 | 0.0078 | 0.17 [0.02, 0.37] | 0.41 |
| | Non-weighted | 1.52 | 0.0060 | 0.18 [0.02, 0.37] | 0.42 |
| Drift rate $v$ (previously encoded) ~ trace persistence $\beta$ | $H_3R$-weighted | 0.02 | 0.0184 | 0.11 [0.00, 0.28] | 0.33 |
| | Non-weighted | 0.02 | 0.0226 | 0.10 [0.00, 0.28] | 0.32 |
| Decision policy $a$ (previously encoded) ~ trace persistence $\beta$ | $H_3R$-weighted | -0.0078 | 0.0033 | 0.19 [0.04, 0.38] | -0.44 |
| | Non-weighted | -0.0077 | 0.0042 | 0.19 [0.03, 0.37] | -0.43 |
| Drift rate $v$ ~ Increasing Complexity $\beta$ (*n*-back) | $H_3R$-weighted | 0.004 | 0.0335 | 0.09 [0.00, 0.26] | 0.30 |
| | Non-weighted | 0.006 | 0.0629 | 0.07 [0.00, 0.23] | 0.26 |

**Note**: Analyses were guided by group-level effects reported in the primary analyses, for both fMRI analyses (clusters) and behavioural/computational data. [1] Bonferroni-Holm correction to adjust for multiple comparisons. [2] $H_3R$-weighted and unweighted connectivity values were compared with $H_3R$-weighted unweighted cluster values, respectively.

**Supplementary Table 10.** Side effects profile during acute administration period (pitolisant vs. placebo) – descriptive statistics and inferential analysis

| *Side effect domain* | Pitolisant (*n*=28) M (S.D.) | Placebo (*n*=29) M (S.D.) | *F*-statistic [df] | *p* |
|---|---|---|---|---|
| **Agitation** | | | | |
| Baseline, M (S.D.) | 0.00 (0.00) | 0.00 (0.00) | -- | -- |
| Follow-up, M (S.D.) | 0.21 (0.50) | 0.21 (0.56) | < 0.00 [1, 54] | 0.958 |
| **Anxiety** | | | | |
| Baseline, M (S.D.) | 0.21 (0.42) | 0.21 (0.42) | -- | -- |
| Follow-up, M (S.D.) | 0.14 (0.36) | 0.14 (0.44) | < 0.00 [1,54] | 0.961 |
| **Appetite, decreased** | | | | |
| Baseline, M (S.D.) | 0.32 (0.72) | 0.11 (0.31) | -- | -- |
| Follow-up, M (S.D.) | 0.29 (0.66) | 0.17 (0.60) | 0.83 [1,54] | 0.367 |
| **Appetite, increased** | | | | |
| Baseline, M (S.D.) | 0.07 (0.26) | 0.17 (0.26) | -- | -- |
| Follow-up, M (S.D.) | 1.00 (1.05) | 0.97 (0.94) | 0.017 [1,54] | 0.898 |
| **Diarrhoea** | | | | |
| Baseline, M (S.D.) | 0.00 (0.00) | 0.00 (0.00) | -- | -- |
| Follow-up, M (S.D.) | 0.00 (0.00) | 0.00 (0.00) | 0.00 [1,54] | 1.000 |
| **Drowsiness/Fatigue** | | | | |
| Baseline, M (S.D.) | 0.14 (0.36) | 0.17 (0.38) | -- | -- |
| Follow-up, M (S.D.) | 1.32 (0.91) | 1.25 (0.84) | 0.24 [1,54] | 0.629 |
| **Dry Mouth** | | | | |
| Baseline, M (S.D.) | 0.14 (0.36) | 0.10 (0.31) | -- | -- |
| Follow-up, M (S.D.) | 0.54 (0.79) | 0.24 (0.51) | 2.577 [1,54] | 0.102 |
| **Indigestion** | | | | |
| Baseline, M (S.D.) | 0.00 (0.00) | 0.00 (0.00) | -- | -- |
| Follow-up, M (S.D.) | 0.00 (0.00) | 0.03 (0.19) | 0.97 [1,54] | 0.330 |
| **Nausea** | | | | |
| Baseline, M (S.D.) | 0.00 (0.00) | 0.00 (0.00) | -- | -- |
| Follow-up, M (S.D.) | 0.18 (0.48) | 0.07 (0.26) | 1.18 [1,54] | 0.282 |
| **Sweating** | | | | |
| Baseline, M (S.D.) | 0.18 (0.39) | 0.10 (0.31) | -- | -- |
| Follow-up, M (S.D.) | 0.14 (0.45) | 0.03 (0.19) | 1.59 [1,54] | 0.213 |
| **Tremors** | | | | |
| Baseline, M (S.D.) | 0.07 (0.26) | 0.00 (0.00) | -- | -- |
| Follow-up, M (S.D.) | 0.04 (0.19) | 0.00 (0.00) | 1.96 [1, 54] | 0.167 |
| **Upset stomach** | | | | |
| Baseline, M (S.D.) | 0.00 (0.00) | 0.00 (0.00) | -- | -- |
| Follow-up, M (S.D.) | 0.00 (0.00) | 0.03 (0.19) | 1.00 [1,54] | 0.330 |
| **Side Effects Aggregate** | | | | |
| Baseline, M (S.D.) | 1.21 (1.37) | 0.90 (1.18) | -- | -- |
| Follow-up, M (S.D.) | 3.86 (2.62) | 3.17 (2.80) | 0.94 [1,54] | 0.337 |

[a] Inferential analysis via baseline-adjusted ANCOVA model across allocation groups (active vs placebo).
**Note:** Table contains data for *N*=57 individuals (114 observations total). *N*=1 individuals were excluded from analyses due to data missingness (baseline).

**Supplementary Table 11.** Self-rated state anxiety, anhedonia and affect across allocation groups

| | Pitolisant (*n*=28) M (S.D.) | Placebo (*n*=29) M (S.D.) | Inferential analysis [a] | |
|---|---|---|---|---|
| | | | *F*-statistic [df] | *p* |
| Dimensional Anhedonia Rating Scale | | | | |
| Baseline, M (S.D.) | 59.07 (4.97) | 57.41 (7.53) | -- | -- |
| Follow-up, M (S.D.) | 58.21 (8.54) | 55.79 (8.60) | 1.83 [1,54] | 0.182 |
| Positive and Negative Affect Schedule | | | | |
| Negative items, baseline, M (S.D.) | 11.68 (3.24) | 10.90 (1.35) | -- | -- |
| Negative items, follow-up, M (S.D.) | 12.89 (3.46) | 11.90 (3.58) | 1.43 [1,54] | 0.238 |
| Positive items, baseline, M (S.D.) | 33.32 (6.63) | 33.03 (7.84) | -- | -- |
| Positive items, follow-up, M (S.D.) | 33.00 (7.14) | 30.86 (8.37) | 2.11 [1,54] | 0.152 |
| Spielberger State Anxiety Subscale | | | | |
| Baseline, M (S.D.) | 27.50 (5.41) | 27.90 (6.14) | -- | -- |
| Follow-up, M (S.D.) | 29.82 (7.59) | 29.31 (9.29) | 0.08 [1,54] | 0.777 |
| Visual Analogue Scale | | | | |
| Negative items, baseline, M (S.D.) | 143.46 (85.36) | 123.76 (82.30) | -- | -- |
| Negative items, follow-up, M (S.D.) | 220.50 (131.00) | 225.28 (125.90) | 0.03 [1,54] | 0.872 |
| Positive items, baseline, M (S.D.) | 725.79 (98.20) | 744.48 (97.45) | -- | -- |
| Positive items, follow-up, M (S.D.) | 630.79 (180.19) | 647.17 (142.01) | 0.20 [1,54] | 0.658 |

[a] Inferential analysis via baseline-adjusted ANCOVA modelling across allocation groups (active vs placebo). **Note:** Table contains data for *N*=57 individuals (114 observations total). *N*=1 individuals were excluded from analyses due to data missingness (baseline).

**Supplementary Table 12.** Mixed-effects linear modelling of subjective cognition throughout the acute assessment period

| | Model estimate [a] | 95% CI | *t* value | *p* |
|---|---|---|---|---|
| Motivation subscale | | | | |
| Time | 0.88 | -0.02, 1.79 | 1.92 | 0.056 |
| Allocation | 2.56 | -2.17, 7.28 | 1.06 | 0.291 |
| Time × allocation | -0.67 | -1.93, 0.59 | -1.04 | 0.301 |
| Mood subscale | | | | |
| Time | -1.55 | -2.40, -0.69 | -3.55 | 0.0005 |
| Allocation | 1.82 | -2.09, 5.74 | 0.91 | 0.363 |
| Time × allocation | -0.34 | -1.53, 0.85 | -0.56 | 0.577 |
| Executive function subscale | | | | |
| Time | -0.78 | -2.69, 1.13 | -0.80 | 0.427 |
| Allocation | 5.30 | -5.61, 16.21 | 0.95 | 0.343 |
| Time × allocation | -3.48 | -6.15, -0.81 | -2.55 | 0.0117 [b] |
| Selective attention subscale | | | | |
| Time | -2.94 | -5.00, -0.88 | -2.80 | 0.006 |
| Allocation | 3.26 | -6.88, 13.40 | 0.63 | 0.529 |
| Time × allocation | -1.71 | -4.59, 1.17 | -1.16 | 0.247 |
| Memory subscale | | | | |
| Time | -3.46 | -5.37, -1.56 | -3.57 | 0.0005 |
| Allocation | -1.45 | -11.42, 8.50 | -0.29 | 0.775 |
| Time × allocation | -1.70 | -4.36, 0.96 | -1.25 | 0.212 |
| Subjective cognition aggregate | | | | |
| Time | -2.98 | -4.49, -1.46 | -3.85 | 0.0002 |
| Allocation | 3.19 | -5.22, 11.60 | 0.74 | 0.459 |
| Time × allocation | -1.66 | -3.78, 0.46 | -1.54 | 0.126 |

[a] Main effect of group (active vs placebo), time, and group × time interactions via time-adjusted mixed-effects linear models with restricted maximum likelihood estimation. [b] While a significant time × group interaction was observed, planned comparisons EMMs showed that allocation groups did not significantly differ at any time point during the testing period ($T_1$ EMM = -2.82 ± 5.14, $p$ = 0.585; $T_2$ EMM = 3.97 ± 5.16, $p$ = 0.443; $T_3$ EMM = 2.83 ± 5.43, $p$ = 0.604; $T_4$ EMM = 8.95 ± 5.14, $p$ = 0.085). **Note:** The SSCS was collected at multiple time points (baseline [$T_1$], pre-scan [$T_2$], post-scan [$T_3$], and visit end [$T_4$]) all of which were included in the model. Table contains data for $N$=57 individuals (114 observations total). $N$=1 individuals were excluded from analyses due to data missingness (baseline).
